# Coral larvae suppress the heat stress response during the onset of symbiosis thereby decreasing their odds of survival

**DOI:** 10.1101/2021.12.10.472165

**Authors:** Sheila A. Kitchen, Duo Jiang, Saki Harii, Noriyuki Satoh, Virginia M. Weis, Chuya Shinzato

**Affiliations:** Department of Integrative Biology, Oregon State University, Corvallis, OR; Statistics Department, Oregon State University, Corvallis, OR; Tropical Biosphere Research Center, University of the Ryukyus, Okinawa, Japan; Marine Genomics Unit, Okinawa Institute of Science and Technology Graduate University, Okinawa, Japan; Okinawa Institute of Science and Technology Graduate University, Okinawa, Japan

**Keywords:** symbiosis onset, coral larvae, heat stress, cell cycle arrest, inflammation, cellular senescence

## Abstract

The endosymbiosis between most corals and their photosynthetic dinoflagellate partners begins early in the host life history, when corals are larvae or juvenile polyps. The capacity of coral larvae to buffer climate-induced stress while in the process of symbiont acquisition could come with physiological trade-offs that alter larval behavior, development, settlement and survivorship. Here we examined the joint effects of thermal stress and symbiosis onset on colonization dynamics, survival, metamorphosis and host gene expression of *Acropora digitifera* larvae. We found that thermal stress decreased symbiont colonization of hosts by 50% and symbiont density by 98.5% over two weeks. Temperature and colonization also influenced larval survival and metamorphosis in an additive manner, where colonized larvae fared worse or prematurely metamorphosed more often than non-colonized larvae under thermal stress. Transcriptomic responses to colonization and thermal stress treatments were largely independent, while the interaction of these treatments revealed contrasting expression profiles of genes that function in the stress response, immunity, inflammation and cell cycle regulation. The combined treatment either canceled or lowered the magnitude of expression of heat-stress responsive genes in the presence of symbionts, revealing a physiological cost to acquiring symbionts at the larval stage with elevated temperatures. In addition, host immune suppression, a hallmark of symbiosis onset under ambient temperature, turned to immune activation under heat stress. Thus, by integrating the physical environment and biotic pressures that mediate pre-settlement event in corals, our results suggest that colonization may hinder larval survival and recruitment creating isolated, genetically similar populations under projected climate scenarios.

## 1. Introduction

The majority of juvenile corals acquire their photosynthetic dinoflagellate symbionts (family Symbiodiniaceae (LaJeunesse et al., 2018)) through horizontal transmission from the surrounding environment (Babcock et al., 1986; Richmond, 1990). Symbiont selection by corals is influenced by multiple factors including host and symbiont genetic backgrounds, algal physiology and environmental availability (Abrego et al., 2009; Coffroth et al., 2001; Cumbo et al., 2013; Little et al., 2004; Quigley et al., 2017; Yamashita et al., 2018). Symbiont colonization of coral larvae may enhance survivorship and extend settlement competency periods, both of which play an important role in larval dispersal (Chamberland et al., 2017; Harii et al., 2009; Suzuki et al., 2013; van Oppen et al., 2001). However, changes in the environment may alter the availability of symbionts, mechanisms controlling symbiont acquisition, or make combinations of some host and symbiont pairings less favorable (Abrego et al., 2012; Cunning et al., 2015; Howe-Kerr et al., 2020; Winkler et al., 2015).

Coral reefs are facing unprecedented thermal stress events globally. Responses of coral larvae to temperature-induced stress are highly variable, depending on the duration of the stress event (from hours to months) and the host symbiotic state. These responses include changes in larval behavior, development, settlement and survivorship with more dramatic changes being observed in symbiotic compared to symbiont-free (aposymbiotic) larvae (Baird, 2006; Jiang et al., 2021; Munday et al., 2009; Nesa et al., 2012; Randall et al., 2009a; Randall et al., 2009b; Schnitzler et al., 2011; Winkler et al., 2015; Yakovleva et al., 2009). For example, elevated temperature impaired symbiosis establishment and greatly decreased larval survivorship in the corals *Fungia scutaria* (Schnitzler et al., 2011) and *Platygyra daedalea* (Jiang et al., 2021). Consequently, as sea surface temperatures continue to rise (Stocker et al., 2013) and time intervals shorten between bleaching events (Hughes et al., 2019), physiologically compromised larvae could result in reduced larval dispersal and recruitment, thereby hindering the development of healthy coral reefs.

To counter extreme stress, corals mount a stress response typified by rapid changes in expression of transcripts related to growth, oxidative stress, protein homeostasis and immunity (Cziesielski et al., 2019; Dixon et al., 2020). Consistent with the coral stress response, the heat-stress response of aposymbiotic larvae is characterized by transcriptional change in heat shock proteins, oxidoreductase activity and cell death (Dixon et al., 2015; Meyer et al., 2011; Polato et al., 2013; Portune et al., 2010; Rodriguez-Lanetty et al., 2009). In contrast, under ambient temperature, onset of symbiosis in larvae by compatible symbiont species elicits little transcriptomic change in the host (Mohamed et al., 2016; Mohamed et al., 2020; Schnitzler et al., 2010; Voolstra et al., 2009; Yoshioka et al., 2021; Yuyama et al., 2018). The transcriptional patterns documented suggest that acquiring symbionts suppresses host immunity and arrests phagosomal maturation of the host-derived symbiosome housing the symbionts (Mohamed et al., 2020; Weis, 2019). Given that symbiotic state plays a role in determining larval health and survivorship (Hartmann et al., 2017), it is imperative to also address transcriptional changes in symbiotic larvae under thermal stress where symbiosis onset may become stressful and could prevent successful symbiosis establishment.

In this study, we simultaneously exposed coral larvae to elevated temperature and colonizing algae to examine symbiont colonization success, larval survival, metamorphosis and host gene expression. We used a factorial design to investigate the interaction of colonization-by-temperature on larvae from the broadcast spawning coral *Acropora digitifera. A. digitifera* is one of the most sensitive species to elevated sea surface temperatures on Okinawan reefs (Loya et al., 2001) where temperatures have remained elevated by a mean of 0.5 °C since 1998 (van Woesik et al., 2011) making it a relevant experimental system to study the molecular responses associated with climate change. Our design allowed us to capture the heat-stress response of aposymbiotic larvae, identify novel transcripts involved in symbiont colonization under ambient conditions, and detect the interactive responses of larvae to the combined treatments that may contribute to their stress response.

## 2. Materials and Methods

### 2.1 Collection and maintenance of coral larvae

Six coral colonies of *A. digitifera* were collected off Sesoko Island, Okinawa Prefecture, Japan (26 °39.1’N, 127°51.3’E) during the 2014 spawning season and placed into ambient temperature (27 °C) seawater tanks. After spawning, buoyant egg and sperm bundles were collected for fertilization, pooled and mixed for 30 min by gentle agitation. Fertilized embryos were washed twice with 0.22 μm filtered seawater (FSW) to remove remaining sperm. Early developmental stages were maintained in ambient temperature FSW with an antibiotic mixture (0.12 mg ampicillin, 0.036 mg kanamycin, 0.007 mg streptomycin) (FSWA) and constant gentle agitation for six days.

### 2.2 Symbiodinium tridacnidorum culture conditions

We inoculated larvae with *Symbiodinium tridacnidorum* sp. nov. (CCMP2465, ITS2 type A3; Lee et al. (2015)) that are acquired readily by early-life stages of *A. digitifera* (Suwa et al., 2010; Yuyama et al., 2005). Cultures of monoclonal *S. tridacnidorum* sp. nov. were grown in f/2 media at 25 °C under a 12h light: 12h dark cycle.

### 2.3 Experimental design

One day prior to starting the experiment, larvae were moved to their respective treatment groups as indicated in Table 1 and placed in a multi-thermo Eyela MTI-201 incubator (Tokyo Rikakikai Co., Japan) at control temperature (26.69 ± 0.18 °C, hourly mean ± stdev) and light intensity of 128.96 ± 32.45 lumens/ft^2^ (Figure S1). Temperature and light intensity of each chamber within the incubator was captured by HOBO Pendant data loggers (Onset Computer Co., MA, USA) submerged in water. Larvae were placed in six replicate 50 ml tubes (TPP, Switzerland) at a concentration of 6 larvae ml^-1^ FSWA with 300 larvae for RNA extraction, 300 larvae for symbiont density quantification and 50 larvae for survivorship analysis.

**Table 1.** Factorial treatment groups. A= aposymbiotic larvae; S= symbiont inoculated larvae, C= contr<u>ol temperature; H= high temperature</u>

| Treatment Group | AC | SC | AH | SH |
| --- | --- | --- | --- | --- |
| Elevated temperature (32 °C) | – | – | + | + |
| Colonization | – | + | – | + |

At six days post-fertilization, larvae, *S. tridacnidorum* and homogenized *Artemia*, a known feeding stimulant in *A. digitifera* larvae (Harii et al., 2009), were pre-incubated for 1 h in either control or elevated (31.98 ± 0.54 °C) temperature. Following the pre-incubation period, 9 × 10^4^ cells ml^-1^ *S. tridacnidorum* were added to the colonization treatments and homogenized *Artemia* was added at a 1:100 v:v to all treatment groups (Table 1). All groups were washed daily with FSWA and incubated at their respective treatment temperatures for the duration of the experiment. The light cycle caused approximately 0.5 °C oscillation in the elevated temperature chamber (Figure S1).

### 2.4 Quantification of colonization success and algal density

Colonization success was determined at multiple time points (3, 6, 12 h post-inoculation (hpi) and 1, 2, 3, 7, and 14 dpi) from a subset of 20 larvae in SC (symbiont inoculated larvae at control temperature) and SH (symbiont inoculated larvae at high temperature) treatments (n= 120 larvae total). Larval squashes were processed by placing 3-4 larvae on a microscope slide and gently pressing down on a coverslip with a pencil eraser. Green fluorescent protein (GFP) of the larvae and chlorophyll autofluorescence of the symbionts were captured by Leica DFC310 FX camera (Leica, IL, USA) using a fluorescent Leica M205 FA stereo-microscope (Leica, IL, USA) with a GFP long pass emission filter. Symbiont presence was tallied and analyzed using a binomial generalized mixed effects model (GLMM) with two fixed factors, temperature and time, and a random factor of replicate tube. Algal density at each time point was quantified as the number of symbionts in 20 randomly sampled larvae of the colonized set using ImageJ v1.47 software (Rasband, 1997-2015). Symbiont counts were analyzed with a GLMM using the same factors and random effect as colonization success but fit to a quasi-Poisson distribution to account for over-dispersion.

### 2.5 Monitoring survival

Survival time was calculated for subsets of 50 larvae (n= 6 different larval populations) between 12 hpi and 14 dpi visually because larvae quickly dissolve after death (Richmond, 1990). Metamorphosis (floating or settled polyps) was noted when present. Larvae that metamorphosed or remained alive at the end of the observation period were excluded from the statistical analysis. Survival time varied significantly across tubes (Figure S2), therefore, between-tube variability was accounted for as a random factor by stratification using a Weibull proportional hazard model in the survival R package (Therneau, 2013). In addition, a binomial GLMM was used to predict the ability or inability of larvae from the different treatment groups to survive beyond the experiment.

### 2.5 RNASeq library preparation and sequencing

At 1 and 3 dpi, approximately 300 treated larvae from each replicate tube (n= 6) of the four treatment conditions were concentrated, snap-frozen and stored at -80°C. At 3 dpi, a subset of 20 individuals from each replicate was randomly removed and scored visually for developmental stage (planula or polyp-like) prior to freezing. Percent metamorphosis was compared using a logistic regression with fixed factors of temperature and colonization.

Total RNA was extracted by tissue homogenization in TRIzol (Invitrogen, CA, USA) with the Polytron P1200 homogenizer (Kinematica, NY, USA), followed by the RNeasy extraction kit (Qiagen, CA, USA) with modifications described by Poole et al. (2016). RNA integrity was assessed using the Agilent 2100 Bioanalyzer (Agilent Technologies, CA, USA) and quantity using the NanoDrop 1000 spectrophotometer (NanoDrop Products, DE, USA). Paired-end sequencing libraries were prepared with 2 μg of total RNA using the TruSeq RNA kit v2 with polyA selection (Illumina, CA, USA) and sequenced on the Illumina Genome Analyzer IIx (Illumina, CA, USA) by the DNA Sequencing Section at Okinawa Institute of Science and Technology Graduate University (Okinawa, Japan).

### 2.6 Gene expression, clustering and network analysis

Short 134 bp paired-end reads were checked for quality with FastQC v0.11.1 (Andrews, 2010) and adaptors removed with cutadapt v1.6 (Martin, 2011). The filtered sequences were mapped against the *A. digitifera* genome assembly (adi_v0.9.scaffold) (Shinzato et al., 2011) and *S. tridacnidorium* genome assembly (symA3_37) (Shoguchi et al., 2018) using Bowtie v2.2.3.0 (Langmead et al., 2012) and TopHat v2.0.12 (Trapnell et al., 2009). Read counts were exported using bedtools v2.21.0 (Quinlan et al., 2010) and differentially expressed genes (DEGs) were calculated using DESeq2 v1.24.0 (Love et al., 2014) fitting the model:

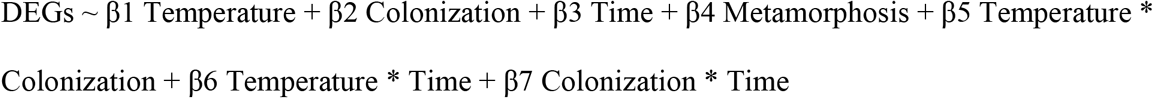

Additional contrasts were constructed to identify DEGs for the SC and AH (aposymbiotic larvae at high temperature) treatment groups on the separate days. The model was run twice to identify transcripts associated with metamorphosis, first treated as a factor (yes or no), and second as a continuous covariate based on the proportion of floating polyps in each sample (Figure S3). Significant DEGs with metamorphosis from both models were removed from subsequent analyses. Hierarchical clustering of the difference in regularized log2 (rlog) counts from the AC (aposymbiotic larvae at control temperature) sample was performed with *hclust* in R (RCoreTeam, 2020). DEGs from at least on treatment (1 dpi= 5368, 3 dpi =3352) were classified based on the highest correlation of rlog counts to a predefined combination of treatment responses as described by Rasmussen et al. (2013). Correlations with significant p-values are reported. Differentiation of transcriptomic profiles among the samples was calculated using a permutational multivariate analysis of variance (PERMANOVA) test (*adonis2* function in vegan R package (Oksanen et al., 2013), 10,000 permutations) for the treatments, sampling times, and interactions as predictors.

Because the *A. digitifera* genome assembly was improved after our data collection, gene models from the adi_v0.9.scaffold genome assembly were compared to the current genome version on NCBI (GCF_000222465.1, version 01-15-2016) using blastp v2.6.0 (-evalue 1e-10, - max_hsps 1 ; (Altschul et al., 1997)). Gene ontology (GO) terms were assigned based on a blastp search against SwissProt and Trembl UniProt databases (release January 31, 2018). A custom database of GO terms was created with *makeOrgPackage* function in AnnotationForge v1.26.0 (Carlson et al., 2019). KEGG orthology was combined from the KEGG Automatic Annotation Server Ver. 1.6a (accessed 2014; https://www.genome.jp/kegg/kaas/) and genome annotation on KEGG API (http://rest.kegg.jp/link/ko/adf). Overrepresentation analyses of GO terms and KEGG orthology terms were tested using *enrichGO* and *enrichKEGG* functions in clusterProfiler v3.12.0 (Yu et al., 2012). P-values were adjusted for multiple comparisons using a Benjamini-Hochberg correction. Z-scores of expression were calculated using GOplot (Walter et al., 2015).

Enrichment of KOG (euKaryotic Orthologous Groups) functional classes based on log_2_ fold-change was performed using Mann–Whitney U tests in the R package KOGMWU (Dixon et al., 2015). The difference in the mean rank for a given KOG class from the mean rank for all other genes, or delta rank, provides a summarized functional response of up- or downregulated genes. Functional similarity was tested between our treatment groups and different growth states (proliferating, quiescent and senescent) of IMR90 human fibroblast (NCBI Gene Expression Omnibus accession: GSE19899, (Chicas et al., 2010)). DEGs were calculated with pairwise comparisons of cell types (e.g. proliferating vs. quiescent) for the human fibroblasts study using the GEO2R NCBI utility. KOG classes were assigned for the human genome (GRCh38.p13) and *A. digitifera* genome using eggNOG-mapper v2.1.5, database 5.0 (Huerta-Cepas et al., 2017).

DEGs were also clustered using a signed, weighted gene co-expression network analysis using WGCNA v1.46 (Langfelder et al., 2008a). A biweight midcorrelation matrix calculated from the variance stabilized counts using DESeq2 (Love et al., 2014) was transformed with a power function of 16. Modules were assigned through unsupervised hierarchical clustering and refined with dynamic tree cutting with a limit of 15 genes per module (Langfelder et al., 2008b).

Hub genes were defined based on intramodular connectivity (kME) > 0.9. GO and KEGG overrepresentation tests described above were performed on each module. Enrichment of DNA motifs were screened as previously reported (Ksouri et al., 2021) at three intervals between - 1500 to 200 bp of the transcript start site (TSS) within each highlighted module against all gene models using the differential RSAT *peak-motifs* tool (http://rsat.sb-roscoff.fr/)(Thomas-Chollier et al., 2012). Up to eight candidate motifs were returned for both the oligo and dyad analysis (Defrance et al., 2008) and candidate motifs were annotated using the footprintDB collection with normalized correlation (Ncor) > 0.4 (Sebastian et al., 2014).

## 3. Results

### 3.1 Symbiont colonization and density

In adult *A. digitifera*, the dominant symbiont is a *Cladocopium* sp. (ITS2 type C1). *A. digitifera* larvae, however, can establish symbioses with multiple genera of Symbiodiniaceae in the laboratory (Cumbo et al., 2013; Harii et al., 2009), including cultured *Symbiodinium tridacnidorum* (ITS2 type A3) that are taken up by acroporid larvae (Yuyama et al., 2005). The colonization and density of *S. tridacnidorum* were measured over time by quantifying the chlorophyll autofluorescence in larvae (Figure 1A). We found a significant interaction of temperature and time on the odds of colonization (binomial GLMM, p < 0.001). Initially, the number of colonized larvae was higher in the elevated temperature (SH) group between 6 and 12 hpi compared to the control temperature group (SC) (binomial GLMM, p= 0.001 for 6 h and 12 h; Figure 1B). However, starting at 1 dpi and for the remainder of the experiment, only 51% of SH larvae were colonized whereas 95% of SC larvae were colonized by 14 dpi (Figure 1B). After accounting for time, the odds of colonization in the SH-treated larvae was 0.37 times lower (95% CI 0.20 to 0.68; binomial GLMM, p =0.002), and the average number of algal cells per larva was 0.03 times lower than the SC-treated larvae (quasi-Poisson GLMM, p < 0.001; Figure 1C).

**Figure 1.**
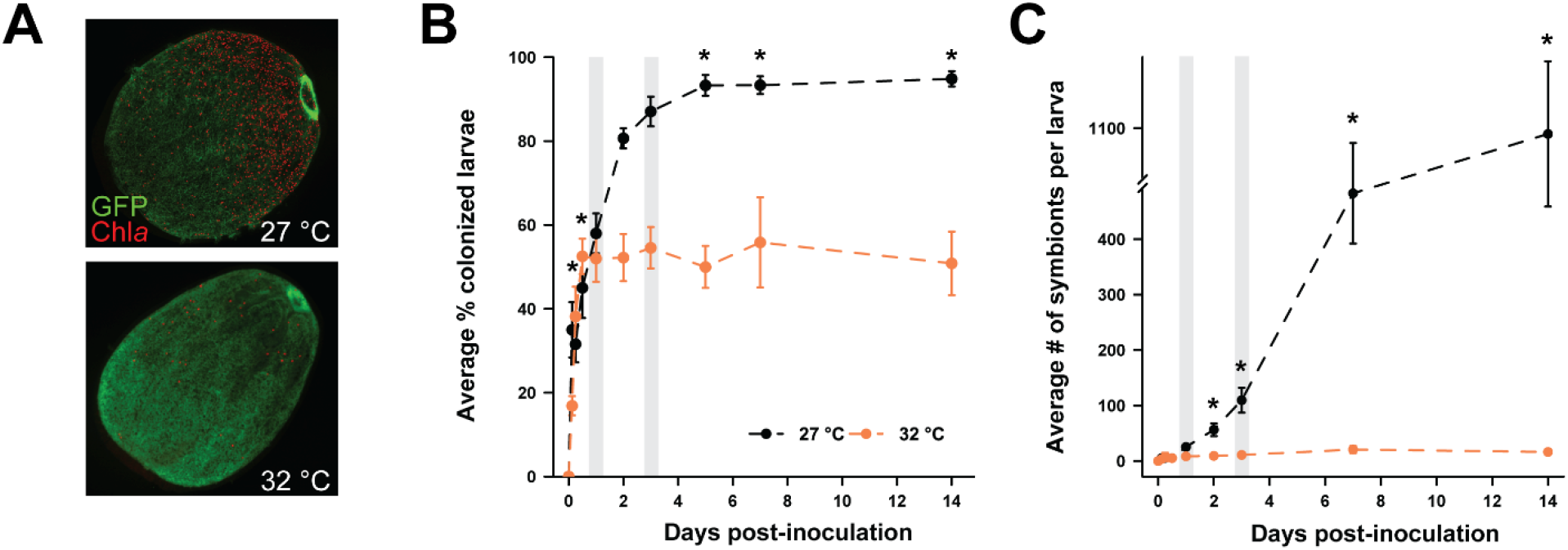
Symbiont colonization and density in *A. digitifera* larvae decreased with elevated temperature. (A) Larvae from 27 °C (black) and 32 °C (orange) treatment groups were inoculated with 9×10^4^ *S. tridacnidorum* at time 0. Chlorophyll (Chl*a)* of the symbionts was used to quantify symbiont colonization (B) and density (C). The green fluorescent protein of the larvae and Chl*a* autofluorescence were both detected with a GFP long pass emission filter. Representative images in A are from 14 dpi in the different temperature treatments. Binomial and quasi-Poisson generalized mixed effects models were fit for colonization success (B) and algal density data (C), respectively. Points represent average estimates with standard error bars. Asterisks denote statistical significance (p ≤ 0.05). RNAseq samples were collected on days one and three (shaded gray bars).

### 3.2 Larval survival

Larval survival was monitored over 14 days with the combined treatment of colonization and elevated temperature. Notably, 84% of the larvae survived the 14 days irrespective of treatment. By the end of the experiment, larvae exposed to the SH treatment had the highest mortality (26.40 ± 6.09 %) while control larvae (aposymbiotic at 27 °C; AC) experienced minimal loss (8.83 ± 1.22%) (Figure 2). There was no significant colonization-by-temperature interaction on survival time (Weibull proportional hazard model, p = 0.46). Nevertheless, elevated temperature (p = 0.003) and colonization (p = 0.040) significantly reduced survival time independently (Figure 2). The probability of larvae surviving beyond 14 days was also significantly affected by both temperature (binomial GLMM, p = 8.0e-4) and colonization (binomial GLMM, p = 0.025), but not by the interaction of these two treatments (binomial GLMM, p = 0.700). After controlling for the effect of colonization, elevated temperature shortened survival time by 90% (Weibull proportional hazard model, 95% CI= 55% to 98%) and the odds of survival beyond 14 days was estimated to be 40% lower than the odds under control temperature (binomial GLMM, 95% CI 24% to 69%). Colonization was associated with a 70% decrease in survival time (Weibull proportional hazard model, 95% CI= 5% to 91%), and the odds of survival beyond 14 days was 55% lower than in the absence of symbionts (binomial GLMM, 95% CI 33% to 93%), after controlling for the temperature effect.

**Figure 2.**
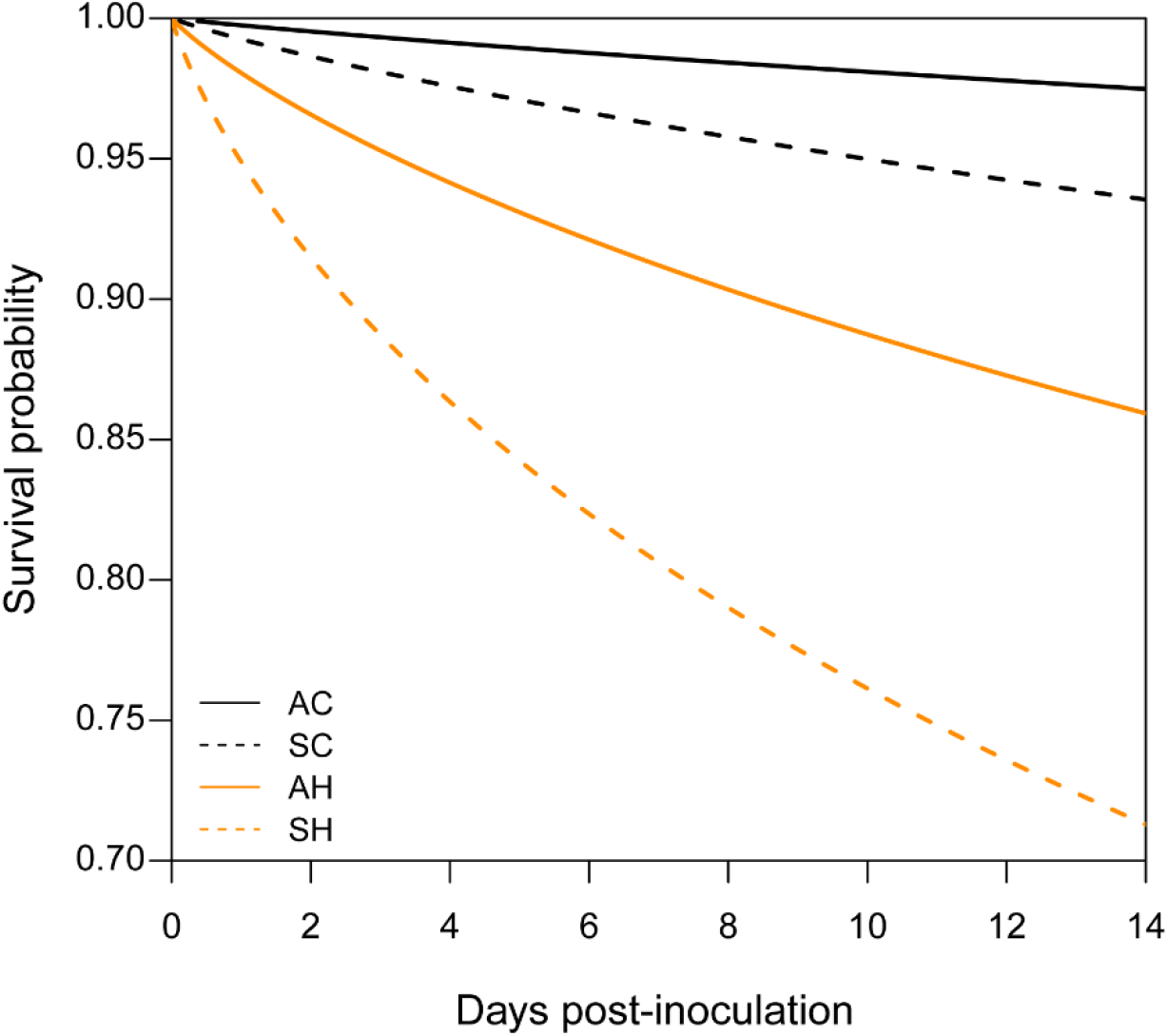
Temperature and colonization decrease survival probability of *A. digitifera* larvae over time. Fitted survival probabilities under the Weibull proportional hazard model for each treatment group (*n*= 300 larvae), averaged across replicate tubes (*n*= 6 tubes per treatment group). The odds of survival decreased significantly with both temperature and symbiosis, but no interaction of these treatments was detected. Solid line = aposymbiotic (A), dashed line = symbiont colonized (S), black = 27 °C control temperature (C), orange = 32 °C high temperature (H).

### 3.3 Metamorphosis observed with elevated temperature and colonization

In addition to the impact of elevated temperature on larval survival and symbiont colonization, it can also affect host development. Studies have noted abnormal larval behavior (Randall et al., 2009a), decreased settlement competence periods (Edmunds et al., 2001; Heyward et al., 2010; Randall et al., 2009b), and atypical floating polyps, i.e. individuals that had undergone metamorphosis without settlement onto a substrate (Putnam et al., 2008), in response to elevated temperature. By 3 dpi, we detected differential proportions of floating polyps in our samples designated for RNA extraction across the treatment conditions (Figure S3). Proportion of metamorphosed individuals ranged from 5% to 85%, resulting in a mixture of both larvae and primary polyp developmental life stages at 3 dpi. While there was no significant colonization-by-temperature effect, both temperature and colonization significantly increased the odds of premature development by 3.31-fold (95% CI= 2.04-5.51) and 5.01-fold (95% CI= 3.02-8.59), respectively (logistic regression, temperature p < 0.001 and colonization p < 0.001).

### 3.4 Gene expression during symbiosis onset with elevated temperature

To capture early transcriptional differences between treatments, we collected samples at 1 dpi and 3 dpi based on the differing larval phenotypes (Figures 1 and 2) and high percentage of metamorphosis (Figure S3). On average 50.43% and 0.018% of the reads mapped to the *A. digitifera* genome assembly and *S. tridacnidorium* genome assembly, respectively (Table S1). The low-mapping rates of the symbiont reads prevented further analysis of the symbiont response.

Given the mixture of life stages present at 3 dpi, we used a principal component analysis to assess the impact of metamorphosis on overall expression (Figure S4A). Metamorphosis accounted for most of the variation along the first principal component (PC1), while the samples from the different temperature treatments separated along PC2. Consistent with these results, the highest proportion of variance in gene expression among the 48 samples was with metamorphosis (15.4%, *F*=13.96, PERMANOVA *p* <0.001) and time (15.3%, *F*=13.88, *p* <0.001) followed by temperature (11.3%, F=10.22, *p*<0.001). To account for the effect of metamorphosis on expression, we included it as a covariate in the statistical model. We recovered 15,211 differentially-expressed genes (DEGs), 8,936 of which were significant with metamorphosis (Table 2 and Figure S4B). These DEGs overlapped with 29% of the DEGs previously identified between larvae and adult *A. digitifera* (Reyes-Bermudez et al., 2016) and were enriched in pathways associated with calcification (upregulation of metallopeptidase activity and extracellular matrix, and downregulation of ion channel activity), and development and growth (upregulation of transcription factor activity and changes in Wnt signaling and Hippo signaling; Figure S5, Tables S3 and S4). Many metamorphosis DEGs were also significant with the heat stress and colonization treatments (62%; Figure S4B). Therefore, we excluded metamorphosis-linked DEGs from further analyses because these patterns could not be separated. With the remaining DEGs, the interaction of colonization-by-temperature explained 2.9% of the variation among samples (F=1.94, PERMANOVA *p*=0.039) while temperature and colonization accounted for 13.4% (F=8.98, p <0.001) and 7.2% (F=4.82, p<0.001), respectively.

**Table 2.**
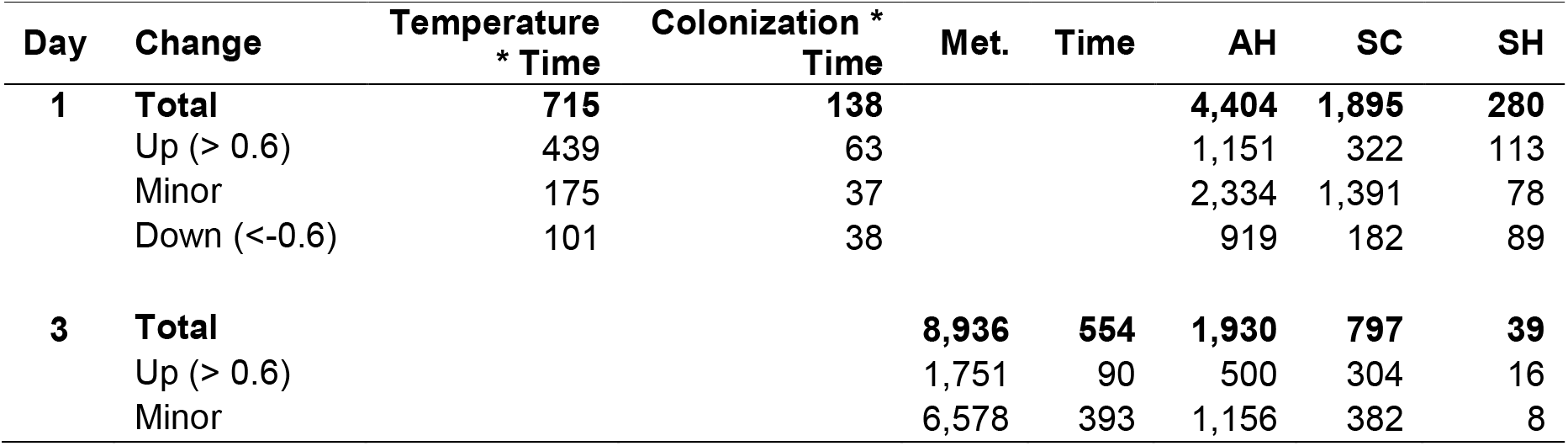
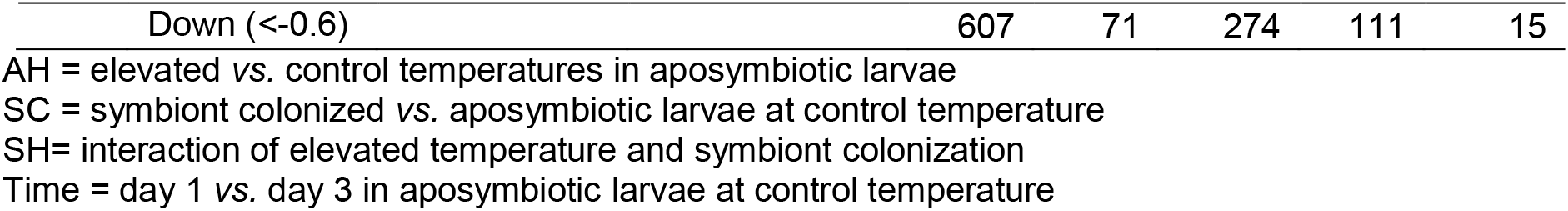
Number of DEGS after excluding those shared with metamorphosis (Met.). Transcripts with expression (log2) < 0.6 and > -0.6 were counted as minor expression.

#### 3.4.1 Responses to the single treatments

Over both sampling days, elevated temperature affected expression more than colonization with a small overlap between treatments (Figure 3A, Table 2 and Table S2). Expression of overlapping DEGs was positively correlated between treatments (SC and AH day 1 Pearson’s correlation= 0.85, p < 0.001; SC and AH day 3 Pearson’s correlation= 0.90, p < 0.001; Figure S6A) and within a treatment across sampling days (SC 1 and 3 dpi Pearson’s correlation= 0.85, p < 0.001; AH 1 and 3 dpi Pearson’s correlation = 0.89, p < 0.001) (Figure S6B). Shared enrichment of GO and KEGG terms for the AH and SC treatments included changes in mitochondrial and proteasome components, and processes such as endocytosis, MAPK signaling, mTOR signaling, and cellular senescence (Table S3 and Table S4). After removing shared terms, transcripts upregulated in RNA processing and cell death, and downregulated in the cell cycle were enriched in AH larvae (Figure S7, and Tables S3 and Table S4). The SC treatment was enriched in transcripts upregulated in intraciliary transport and cytokine receptor binding and downregulated in oxidative phosphorylation (Figure S7, and Table S3 and Table S4).

**Figure 3.**
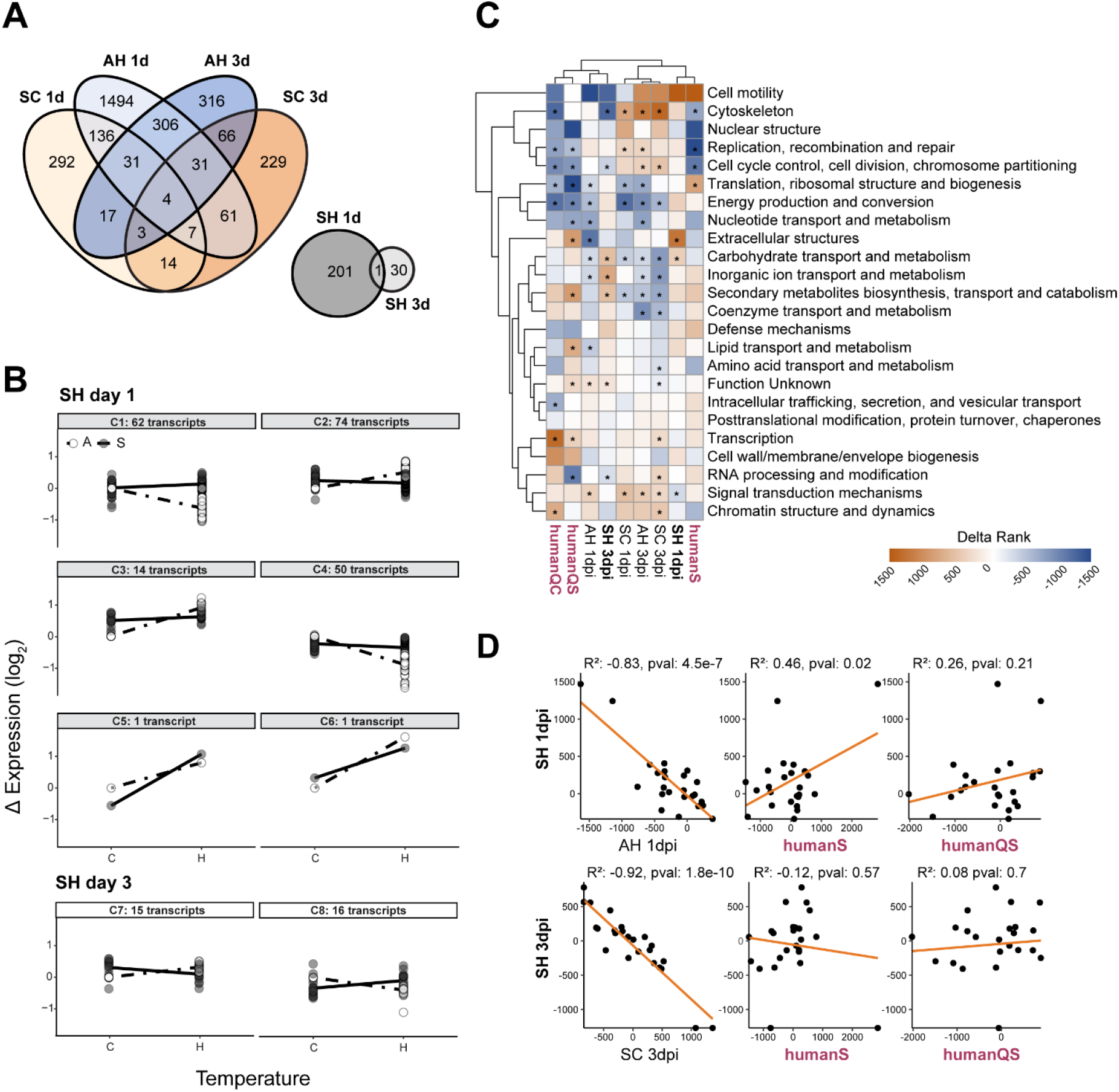
Differentially expressed genes by treatment. (A) Shared DEGs separated by sampling day between AH and SC treatment groups and SH treated-larvae are presented in the Venn diagrams. (B) The interaction of temperature and colonization was visualized with interaction plots for the 232 DEGs in SH-treated larvae. Hierarchical clustering of a Euclidian distance matrix generated from the difference in rlog transformed counts from AC larvae resulted in 8 expression clusters (C1-C8) with the transcripts numbers ranging from 1 to 73. Lines represent the average difference in rlog expression of all transcripts in that cluster whereas points represent individual differences in rlog expression for each transcript for the given condition. Aposymbiotic = ○ and dashed line, symbiont colonized = • and solid line. (C) Heat map of clustered KOG classes and treatment conditions of this study and human fibroblasts in quiescence (QS= serum-induced, QC=contact-induced) or senescence (S) (Chicas et al., 2010). The colors represent the delta-rank of log2 fold-change where positive values (orange) indicate enrichment of up-regulated genes. Asterisks denote significantly enriched KOG classes (FDR=0.05). (D) Pairwise Pearson’s correlation plots of KOG delta-ranks for select treatments.

#### 3.4.2 Responses to the combined treatment

In contrast to the singular responses of SC-and AH-treated larvae, we identified 202 and 31 DEGs that respond differently to the interaction of colonization-by-temperature at 1 and 3 dpi, respectively (Figure 3A). To explore the interactions further, the DEGs were clustered based on fold-change into eight groups (C1-C8) ranging from one to 73 DEGs per cluster (Figure 3B). These interactive effects can result in “*combinatorial*” (response of the combination differs from similar responses of the single treatments), “*prioritized*” (responses of the single treatments differ while the combination response remains at the same level of one of these treatments), or “*canceled*” (responses of the single treatments differ but the combination response returns to control level) responses as described by Rasmussen et al. (2013). Other responses where there is no apparent interaction include “*similar*” (response level is the same for all treatments) and “*independent*” (response of a single treatment and the combined treatment are the same). Using these classifications, two patterns emerged in the expression clusters, 1) 61% of the DEGs in C1 to C4 were classified as *canceled* (i.e. elevated temperature response reduced by the addition of symbionts); and 2) 48% of the DEGs in C7 and C8 were classified as *combinatorial* with similar expression between AH and SC larvae but the opposite response in SH larvae. Consistent with these results, the SH larvae transcriptomes were functionally distinct from the AH and SC transcriptomes on each sampling day based on the KOG delta ranks (Figure 3C). The most significantly upregulated KOG class in SH larvae was ‘Extracellular structures’ on 1 dpi, and ‘Carbohydrate transport and metabolism’ on both days.

To further examine the interaction of colonization-by-temperature, we classified all DEGs that were significant with at least one treatment (Table S2). Most transcripts were classified as non-interactive, either as *independent* (1 dpi=37%, 3 dpi=44%) or *similar* responses (1 dpi= 29%, 3 dpi=33%). Of the interactive responses, the number of *prioritized* colonization responses was 3-fold higher than *prioritized* temperature responses at 1 dpi, suggesting a physiological trade-off of colonization with elevated temperature. Likewise, the number of *canceled* temperature responses was 10-fold higher than *canceled* colonization responses. This resulted in a significant negative correlation of -0.83 and -0.43 between the KOG delta ranks of SH larvae and AH and SC larvae, respectively (only AH shown, Figure 3D). By 3 dpi, after the initial period of symbiont colonization ended, the number of *prioritized* colonization responses and *canceled* temperature transcripts were only a fraction of the equivalent responses, leading to a similar negative correlation between SH larvae and either AH or SC KOG delta ranks, respectively (only SC shown, Figure 3D).

From the different enrichment analyses (GO, KEGG and KOG), we found that both AH and SH larvae shared the downregulation of cell cycle genes (Figure 3C and Table S4). To explore the different cellular states that would lead to the downregulation of cell cycle, we compared the transcriptomic profiles of the different treatments to human fibroblasts in either quiescence, a reversible state of cell cycle arrest, or senescence, a non-reversible, arrested state (Chicas et al., 2010). We found that the transcriptomic response of SH larvae was functionally similar to senescent cells (Pearson’s correlation = 0.46, p=0.02), but not quiescent cells at 1 dpi (Figure 3D), suggestive of permanent cell cycle arrest, whereas the transcriptomic response of AH larvae was not significantly correlated with either cellular state.

### 3.5 Co-expression networks identify additional expression patterns associated with the SH treatment

To identify additional transcripts that may be functionally related or share the same regulatory program as the colonization-by-temperature DEGs, we constructed co-expression networks (Carter et al., 2004). We investigated five modules – M12, M14, M15, M21, and M24 – that were positively correlated with elevated temperature and three modules – M13, M15, and M25 – that were moderately correlated with colonization (Figure 4A).

**Figure 4.**
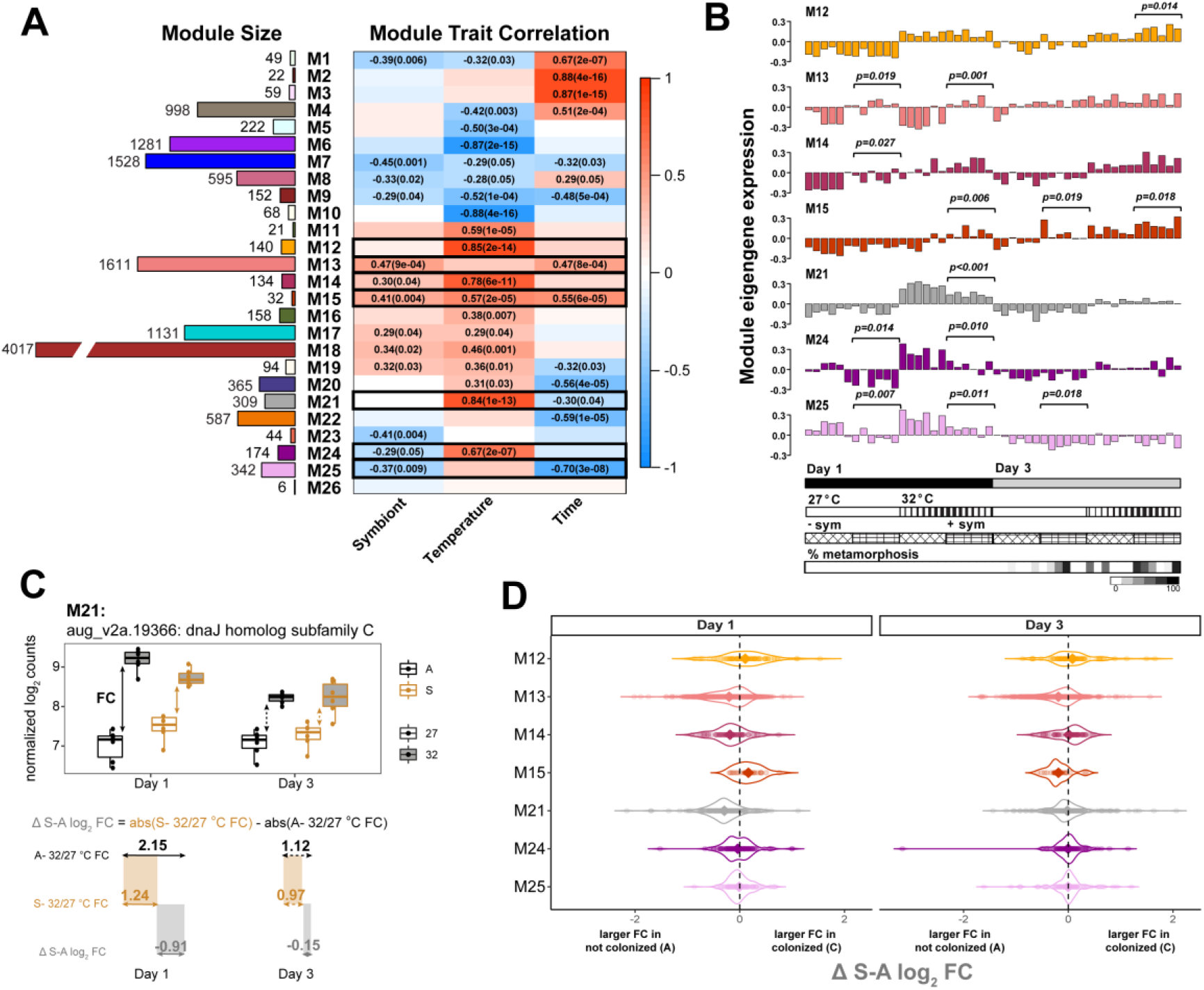
Comparison of co-expression modules correlated to experimental treatments. **(A)** Twenty-six co-expression modules were clustered using an adjacency matrix of variance stabilization transformed counts using WGCNA (Langfelder *et al*., 2008a). Module size, or the total number of transcripts clustered in each module, are represented by the colored bars on the left and the biweight midcorrelation (*p*-value) of each treatment to a given module is presented on the right. The correlation strength is indicated by color from 1 (red) to -1 (blue). (B) For select modules (black boxes), an average sample eigengene value was calculated from the expression of all transcripts in that module. Bars at the bottom correspond to experimental conditions in the bar graphs above (day = black is 1 or gray is 3; temperature = white is 27 or lines is 32 °C; colonization = diamond hatch mark is − sym or grid is + sym; and percent metamorphosis is grayscale). Student’s T-test results are presented above + sym samples that significantly differed with the – sym samples in each respective temperature. (C-D) Effect size of temperature and colonization on expression of module transcripts. In panel (C), the difference in the absolute value of colonized high to low temperature fold-change from the absolute value of the not colonized high to low temperature fold-change (Δ S-A log2 expression) was calculated as exemplified for one of the hub genes in module M21. (D) Violin plots show the distribution of Δ S-A log2 expression for significant transcripts within a module, where individual points represent transcripts and the mean difference is represented by the diamond. Negative values indicate larger fold-changes in the aposymbiotic larvae while positive values indicate larger fold-changes in the symbiont colonized larvae.

Of the modules that were positively correlated with elevated temperature (M12, M21 and M24, Figure 4A and Figures S8-S10), M12 was enriched in transcripts involved in cell death (Table S5) and all modules were overrepresented by KEGG pathways involved in inflammation response and misfolded protein processes (Table S6). To identify potential regulators of the gene networks, DNA sequence 1,500 bp upstream to 200 bp downstream of the transcription start site (TSS) were scanned for shared DNA motifs. Genes in M12 were enriched in motifs annotated as transcription factor Nuclear factor-κB *(NF-κB)* and DNA binding motif RelA, a regulator of NF-κB (Table S7). In M24, genes were enriched in homeobox domains and zinc finger binding sites, but no enriched motifs were found for M21 genes (Table S7).

Eigengene expression of colonized samples in modules correlated with colonization (M13 and M25) was significantly upregulated or downregulated at 1 dpi irrespective of temperature treatment (Student’s T-test; Figure 4B, Figure S11-S12), suggesting that the transcripts might be involved in initial symbiont recognition. There were 12 motifs identified upstream of the TSS of M13 genes, including binding sites for multifunctional transcription factors MafF and YY2, and 10 novel motifs (Table S7). Transcripts in M13 module were associated with endocytosis, cytoskeleton, apoptosis and mTOR signaling (Table S5 and Table S6) and were highly-connected with 59 hub genes (Table S8), whereas transcripts in M25 were associated with the mitochondrial electron transport chain (Table S5).

The last two modules, M14 and M15, were both positively correlated with colonization and elevated temperature (Figure 4A) and their eigengene expression was significantly higher in colonized larvae at different temperatures on 1 dpi (Student’s T-test, Figure 4B). Each module had five hub genes with increasing expression with symbiotic-state and temperature (Table S8, Figures S13-S14). M14 genes were enriched for inhibition of NF-κB and inflammatory response, while M15 genes were enriched for endocytosis and MAPK signaling pathway (Figures S14, and Tables S5 and S6). Enriched regulatory motifs in M14 include binding sites of nuclear respiratory factor 1 (Nrf1) and E2F8 transcription factors, and transcriptional repressor CTCFL, whereas M15 was enriched in binding sites associated with C2H2 type zinc finger domains (Table S7).

While the directional change in expression of transcripts were similar between some treatment groups within a co-expression module, the effect size of the treatments varied. For example, in M21, elevated temperature treatment increased the log_2_ fold-change of *dnaJ homolog subfamily C member 3-like* in both symbiotic states, but the fold-change in symbiotic larvae was 0.91 lower than aposymbiotic larvae at 1 dpi (Figure 4C). The difference in log_2_ fold-change for each DEG within a module was calculated based on either symbiotic-state (Δ S-A log_2_ expression, Figure 4D) or temperature treatment (Δ 32-27 °C log_2_ expression, Figure S19). We found that symbiotic condition lowered the expression of M21 and M13 transcripts relative to aposymbiotic larvae at 1 dpi (Figure 4D) whereas there was an increase in mean Δ log_2_ expression of heat-stressed larvae for M13 only (Figure S15). These differences in the effect size indicate that the combination of treatments lowers the transcriptional response of key genes involved in heat stress response (M21) and negates temperature-induced downregulation of genes involved in colonization (M13). The Δ log_2_ expression of transcripts in modules M14 and M15 represent a set of genes where temperature and colonization effect sizes were shifted in the same direction resulting in either larger fold-changes in non-colonized, control larvae (M14) or larger fold-changes in colonized, heat-treated larvae (M15) that was reversed by 3 dpi (Figure 4D and Figure S15).

## 4. Discussion

By combining symbiosis onset with thermal stress, we revealed complex transcriptional responses of coral larvae that coincided with their decreased symbiont colonization and survival over the two-week experiment (Figure 5A). Like symbiotic anemones (Cleves et al., 2020), the majority of the transcriptomic responses occurred in a single treatment independent of the other, likely as a result of activating different, unrelated processes (Folt et al., 1999). Thermal stress also activated a broader transcriptional response than did colonization, consistent with previous studies on coral larvae (Dixon et al., 2015; Meyer et al., 2011; Mohamed et al., 2016; Mohamed et al., 2020; Polato et al., 2013; Portune et al., 2010; Rodriguez-Lanetty et al., 2009; Schnitzler et al., 2010; Voolstra et al., 2009; Yoshioka et al., 2021; Yuyama et al., 2018). Where transcriptomic responses overlapped on the first day, expression of transcripts important for the heat stress response (HSR) was reduced by symbiont colonization in SH larvae (expression clusters C1-C4, and module M21). Moreover, the SH treatment disrupted expression of transcripts involved in cell proliferation, immunity, and apoptosis resulting in either opposing (expression clusters C7 and C8, and modules M24 and M25) or additive (modules M14 and M15) responses compared to each treatment individually. Taken together, these patterns reveal physiological trade-offs when colonization and thermal stress occurred concomitantly: 1) colonization constrained the host HSR and 2) thermal stress activated a host immune response.

**Figure 5.**
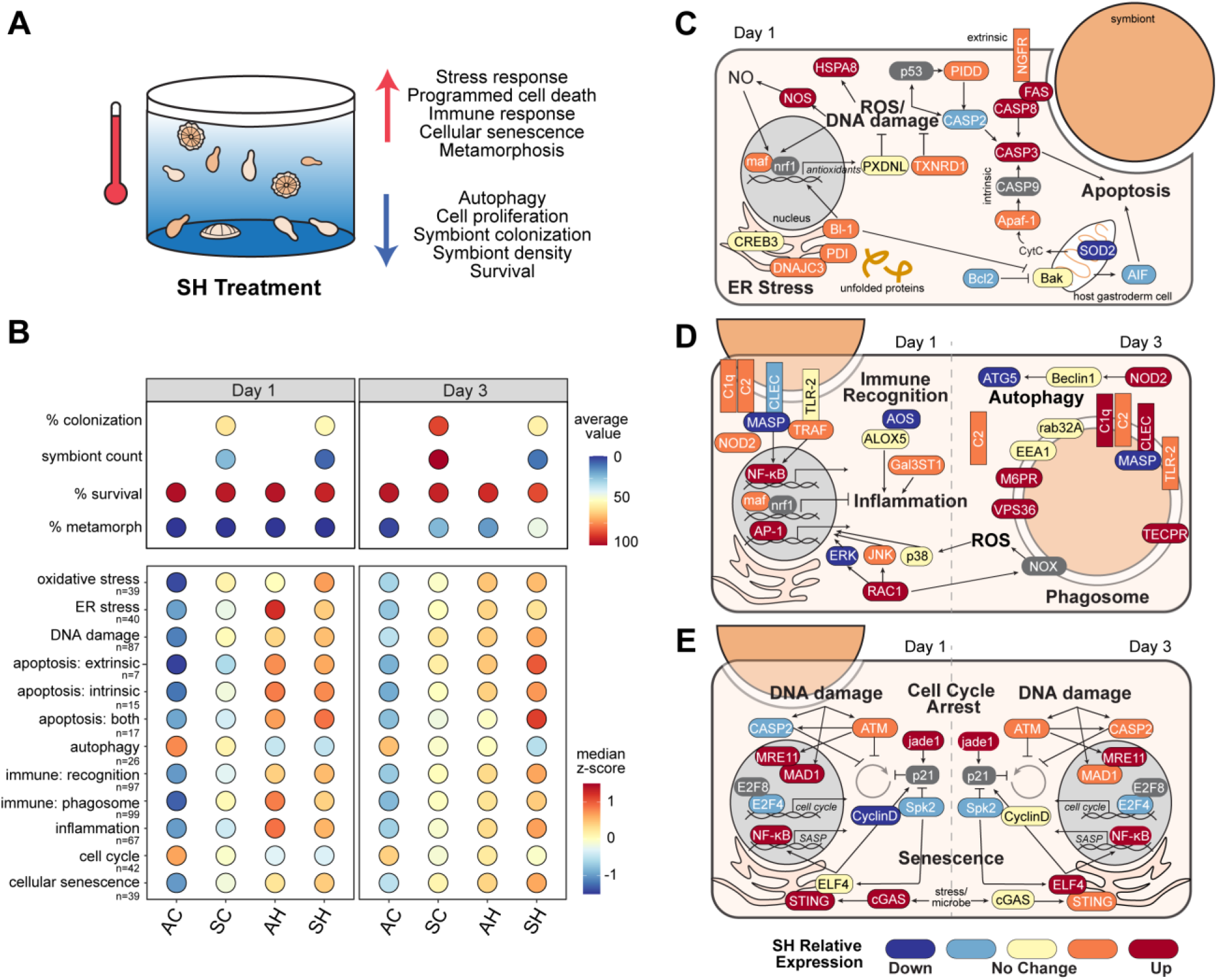
Summary of phenotypic and transcriptomic response of SH-treated larvae. (A) Schematic overview of the changes observed in SH larvae relative to individual treatments. (B, top) Average percent symbiont colonization, algal numbers, percent survival and percent metamorphosis for each treatment group on 1 and 3 dpi. (B, bottom) Median normalized expression (z-score) for all transcripts within a cellular pathway. Expression was transformed to show activation of each respective pathway. The color of the circles represents the average (top) and median (bottom) values for each metric, respectively. (C-E) Cellular processes altered by SH treatment: (C) stress response and cell death on 1 dpi, (D) immune response on 1 and 3 dpi, and (E) cell cycle and senescence on 1 and 3 dpi. The gene color indicates the qualitative expression relative to the other treatments shown in Figure S16, with blues and reds depicting decreased and increased expression in SH animals, respectively. Genes in grey were not differentially expressed or excluded. The genes and gene symbols for each category are listed in Table S9.

### 4.1 Compared to AH larvae, SH larvae fail to mount a full heat-stress response

Thermal stress was the dominant driver of expression differences in the larvae. Under thermal stress, a series of evolutionarily conserved mechanisms to offset the deleterious effects of temperature, known as the HSR, are triggered (Cziesielski et al., 2019; Hofmann et al., 2010). The degree to which thermal compensation is achieved through plastic responses of larvae can be limited by the prior parental environmental history and the presence of multiple stressors (Rivera et al., 2021; Seebacher et al., 2015). We found that the magnitude and/or direction of change in HSR genes differed in SH larvae relative to the stereotypical response of AH larvae (Figure 5B). Specifically, regulation of genes related to oxidative stress, endoplasmic reticulum (ER) stress and apoptosis differed in SH larvae (Figure 5B and S16A).

Oxidative stress can be an indicator of cellular stress and symbiosis dysfunction in cnidarians (Lesser, 1996; Louis et al., 2017). It is characterized by elevated levels of free radicals (reactive oxygen species (ROS) and nitric oxide (NO)) in the cell that activate pro-death pathways. Antioxidants (*PXDNL, TXNRD1, SOD1* and *SOD2*) that scavenge free radicals showed contrasting expression between AH- and SH-treated larvae on 1 dpi (Figure 5C and S16A) but became more similar by 3 dpi (Figure S16A). Antioxidants are commonly upregulated in juvenile corals under thermal stress (Polato et al., 2010; Ritson-Williams et al., 2016; Voolstra et al., 2009; Yakovleva et al., 2009), suggesting that the process of symbiont uptake altered the antioxidant defenses of the host HSR. Moreover, unique combinatorial upregulation of a *nitric oxide synthase* (*NOS*) in the SH treatment on 1 dpi suggests higher levels of cytotoxic NO in these larvae relative to the those in the individual treatments (Figure 5C). Elevated NO, if present, could contribute to cell death (Hawkins et al., 2013) and symbiont loss (Wang et al., 2011).

Oxidative and thermal stress can also lead to disruption of protein homeostasis in the ER, a process regulated by molecular chaperones (Cao et al., 2014). Expression of three molecular chaperones (two *HSPA8* and *PDI*) in AH and SH larvae at 1 dpi (Figure 5C) suggests reduced protein biosynthesis in conjunction with increased protein mis-folding in the temperature-treated larvae, a trigger of the unfolded protein response (UPR). Activation of the UPR attempts to buffer ER stress to restore protein homeostasis, as seen in the transcriptional response of *Acropora hyacinthus* to episodic thermal stress (Ruiz-Jones et al., 2017). However, failure to overcome ER stress results in cell death (Walter et al., 2011). In this study, the ER stress response was reduced in SH relative to AH larvae (Figure 5B) as evidenced specifically by the canceled temperature-induced expression of ER stress responsive transcription factor *cAMP response element-binding protein 3* (*CREB3*), and slight upregulation of ER stress chaperone *DnaJ homolog subfamily C member 3 (DNAJC3)* in SH relative to AC larvae (Figure 5C) (Boriushkin et al., 2014; Kondo et al., 2007). These findings suggest that the severity of ER stress is partially attenuated in SH larvae.

Unresolved cellular stress can lead to the activation of apoptosis and autophagy, two highly regulated, destructive processes that are also proposed mechanisms of symbiont removal in cnidarians (Downs et al., 2009; Dunn et al., 2002; Dunn et al., 2007; Tchernov et al., 2011). We found that controlled death via apoptosis was elevated relative to autophagy in response to heat stress (Figure 5A). Median expression of transcripts involved in apoptosis was similar between AH and SH larvae on 1 dpi (Figure 5B), but a detailed examination revealed differences conditional to colonization (Figure 5C and S16A). Apoptosis can be activated in one of two ways: 1) the mitochondrial-mediated intrinsic pathway, or 2) the damage-activated, receptor-mediated extrinsic pathway (Marino et al., 2014). Evidence of extrinsic apoptosis was observed with elevated temperature irrespective of symbionts on 1 dpi, whereas intrinsic apoptosis was reduced in SH larvae (Figure S16A). For example, *Bcl-2 antagonist killer (BAK)*, a protein that initiates mitochondrial-mediated apoptosis, chaperone *PDI* that induces Bak-dependent mitochondrial permeabilization and *apoptotic protease activating factor 1* (*Apaf-1*) that cleaves procaspase 9 following permeabilization, were upregulated with temperature, but less so in SH compared to AH larvae (Figure 5C and S16A) (Zhao et al., 2015). Consistent with these patterns, ER-bound anti-apoptotic *Bax inhibitor 1* (*Bl-1*) was upregulated in SH larvae (Figure 5C), perhaps to suppress or attenuate intrinsic apoptosis (Ishikawa et al., 2011). By 3 dpi, extrinsic and intrinsic apoptosis remained elevated in SH larvae compared to AH larvae (Figure 5B). This could contribute to the differences in their survival. Taken together, the combined treatment of symbiosis onset and heat stress increased oxidative stress but reduced ER stress, leading to induction of distinct cell death pathways.

### 4.2 Immune activation in SH larvae is evident in contrast to immune suppression in SC larvae

The transcriptomic response of SH larvae also significantly differed from SC larvae on both sampling days such that key immune and inflammatory pathways were activated in SH larvae (Figure 3 and 5B). However, the inflammatory response was reduced in SH larvae relative to AH larvae (Figure 5B). These results support a growing number of studies that show that thermal stress reverses host immune suppression in symbiotic cnidarians and results in phagosomal maturation and inflammation (DeSalvo et al., 2010; Mansfield et al., 2017; Pinzón et al., 2015; Starcevic et al., 2010).

Expression of transcripts putatively involved in symbiont recognition displayed a mixture of canceled (*TLR-2* and *NOD2*) and additive (16 *NOD*-like and two *intelectin* genes) patterns in SH larvae (Figure 5D and S16B). This mixed response could be due to the extensive lineage-specific expansion of TLRs (Poole et al., 2014) and NODs (Hamada et al., 2012) in *A. digitifera* that may participate in both immune and tolerogenic signaling. By 3 dpi, a *C-type lectin* (*CLEC*) was upregulated in SH larvae (Figure 5D). Lectins are carbohydrate-binding proteins on host cells that bind to symbiont glycans (Kvennefors et al., 2010), contributing to their recognition and/or removal. Lectins are also functionally linked to the innate immune complement system through mannose-binding serine proteases (MASP) that may play a role in the microbe recognition (Poole et al., 2016). In contrast to *CLEC* expression, *MASP* was highly expressed in the AC larvae but showed decreased expression in colonized and temperature-stressed larvae (Figure 5D and S16B). However, upregulation of components of a different branch of the complement pathway (*C1q* and *C2*, also known as *factor B*) in addition to increased *CLEC* may therefore enhance removal of symbiotic and/or apoptotic cells through independent activation of phagocytosis (Eddie Ip et al., 2009; Galvan et al., 2012) (Figure 5D). Various phagosome-associated genes (*EEA1, M6PR, VPS36, TECPR1*, and *Rab32*) were also upregulated at some point with temperature, indicative of phagolysosome maturation and possibly symbiont removal in SH larvae (Figure 5B and D).

Another part of innate immunity, inflammation, also plays a role in cellular homeostasis under genotoxic stress (Chovatiya et al., 2014). Reduced inflammation in SH larvae relative to AH larvae was evident by decreased expression of *arachidonate 5-lipoxygenase* (*ALOX*) and *allene oxide synthase* (*AOS*), enzymes active during inflammation in corals (Libro et al., 2013; Ricci et al., 2019) (Figure 5D). Additive expression of genes in SH larvae that negatively regulate NF-κB (module M14, Table S5), a key regulator of the inflammatory and stress response (Liu et al., 2017), and enrichment of binding sites of transcription factor Nrf1, a repressor of inflammation (Biswas et al., 2010), also point to a reduced inflammatory response (Figure 5D). Thus, anti-inflammatory mechanisms activated by the host in the presence of symbionts or by symbiont-derived molecules may counter inflammation as a way to prevent symbiont loss (Erturk-Hasdemir et al., 2019). Nevertheless, activation of stress responsive genes *c-Jun N-terminal kinase* (*JNK*), and *AP-1* in temperature-treated larvae indicates some degree of inflammation in SH larvae not experienced by SC larvae (Figure 5D) (Courtial et al., 2017). Consistent with these patterns, canceled expression on 3 dpi of multiple *TRAFs, galactosylceramide sulfotransferase*, an enzyme that triggers an AP-1 mediated inflammatory response (Jeon et al., 2008) along with *AP-1* (Figure S16C), suggest a moderate inflammatory response of SH larvae throughout early symbiosis.

### 4.3 Heat stress with colonization induces cell cycle arrest and cellular senescence

Unlike SC larvae, colonization and symbiont density plateaued after 1 dpi in the SH larvae, and remained low throughout the experiment at comparable numbers to those observed in *S. tridacnidorum* colonized larvae of *Acropora millepora* held at 32 °C for six days (Cumbo et al., 2018). Elevated ROS and NO levels, as seen in all treated larvae (Figure 5B), can result in DNA damage that triggers cell cycle arrest (Barzilai et al., 2004; Nguyen et al., 1992). Transcriptional evidence of DNA damage and cell cycle arrest was detected in all treatments by 3 dpi (Figure 5B and Figure S16D). Critical genes involved in the cell cycle G1-S transition (*Skp2* and *cyclin D2*), and cell proliferation (transcription factor *E2F4*) were downregulated whereas *jade-1*, an activator of cyclin-dependent kinase inhibitor p21 (Siriwardana et al., 2015), was upregulated in both heat-stressed and colonized larvae, suggesting that a population of host cells was arrested at the G1 phase of the cell cycle (Figure 5E). Moreover, transcripts in module M14, where expression was magnified in SH-treated larvae at 1 dpi, were enriched for binding sites of cell proliferation transcriptional repressor *E2F8* (Logan et al., 2005). By 3 dpi, heat-treated larvae showed evidence of arrested cell proliferation via the DNA damage response (upregulation of DNA repair genes *MRE11* and *MAD1*, and DNA damage sensors *ATM* and *CASP2* (Figure 5E and Table S2)).

Our results for heat-treated larvae are perhaps not surprising given that constant thermal stress arrests cell cycle progression as part of the conserved HSR of eukaryotes (Richter et al., 2010), as observed in the coral *Stylophora pistillata* (Maor-Landaw et al., 2014) and cultured Symbiodiniaceae (Fujise et al., 2018). However, it is surprising that host cells arrested cell proliferation following colonization. Previous studies have shown that host cell proliferation increases in the presence of symbionts, but less so at the larval and metamorphic stages (Lecointe et al., 2016; Tivey et al., 2020). Reports of symbiont contributions to the host energetics during early life stages are mixed with evidence of extra energy reserves in some symbiotic larvae (Harii et al., 2010; Richmond, 1987) while other studies found little to no evidence of contributions from the symbiont (Hartmann et al., 2019; Kopp et al., 2016). Nevertheless, the host may control symbiont growth when resources are scarce or nearing metamorphosis by inducing cellular senescence, a permanently arrested but metabolically active state.

Cellular senescence is a tightly regulated cellular program employed during development, tissue repair, and microbial infections (Kumari et al., 2021; Rhinn et al., 2019; Wei et al., 2018). Components of the cellular senescence pathway were enriched in all treated larvae with a moderate correlation of the SH transcriptomic response to senescent human fibroblasts (Figure 3D, Table S4 and S6). Moreover, the upregulation of *cyclic GMP-AMP synthase* (*cGAS*) with colonization and multiple *stimulator of interferon genes (STING)* with temperature and colonization may promote senescence through secretion of pro-inflammatory factors, referred to as the senescence-associated secretory phenotype (Glück et al., 2017)(Figure 5E). Pro-inflammatory signaling may act to reinforce growth arrest, promote apoptosis resistance in the senescent cells and create an inflammatory microenvironment for senescent cell clearance (Kumari et al., 2021). We hypothesize that larval cells may enter senescence to regulate symbiont growth and/or clear damaged cells to enhance their survival until they settle and undergo metamorphosis. Given the heterogeneity of the senescent phenotype, though, additional cellular markers beyond transcriptome profiles would be needed to confirm the presence of senescent cells within the larvae and whether they are symbiont-containing, gastrodermal cells.

### 4.4 Colonization by symbionts is not beneficial to coral larvae at elevated temperatures

We found that colonization not only reduced larval survival but also accelerated host development at both elevated and ambient temperatures (Figure 5B). The debate remains open on whether the benefits of acquiring symbionts during the larval stage of coral development outweigh the costs under non-stressful conditions (Chamberland et al., 2017; Hartmann et al., 2017; Mohamed et al., 2020). Enhanced survival and dispersal windows may be experienced by some acroporid species (Suzuki et al., 2013), but others, including *A. digitifera* shown here, may experience physiological trade-offs that impair proper development (Nesa et al., 2012). What is generally agreed upon is that elevated temperature increases larval mortality (but see Winkler et al. (2015)). Higher mortality from elevated temperature has been observed in larvae from other Okinawan acroporids including *A. muricata* (Baird, 2006) and *A. intermedia* (Yakovleva et al., 2009); however, only *A. intermedia* larvae showed a difference in survival with colonization.

An unexpected outcome of our treatment conditions was premature metamorphosis of the larvae. Temperature is known to expedite development in corals (Heyward et al., 2010) and similar results have been observed in experiments with symbiotic larvae (Edmunds et al., 2001; Hartmann et al., 2019; Putnam et al., 2008). In the case of *A. digitifera*, metamorphosis is density-dependent, but only in the presence of settlement inducers (Doropoulos et al., 2018). Therefore, given that the larval densities were constant for all treatments and no known settlement inducers were provided, the presence of symbionts may also contribute to the developmental cues that trigger the transition of larvae to primary polyps and could be tested in future studies.

## 5. Conclusion

Our data show that under elevated temperature, *A. digitifera* larval survivorship and *S. tridacnidorum* densities were significantly reduced. Temperature and symbiont colonization also initiated premature metamorphosis. These results suggest that corals face severe challenges in recruitment, survivorship, and therefore fitness with climate change. The transcriptomic snapshots before and after observable phenotypic differences to the single and joint treatments revealed trade-offs in the host physiology. The combination led to the activation of growth arrest, inflammation and DNA damage responses that all point to the emergence of cellular senescence, a protective measure to offset the combined stressors. Overall, these findings expand our understanding of larval response to predicted climatic events.

## Acknowledgements

We would like to thank the students and support staff at Tropical Biosphere Research Center of University of Ryukyus and Okinawa Institute of Science and Technology Graduate University for assistance in coral collection and larval rearing. We thank Dr. Eiichi Shoguchi for providing symbiont cultures and Maria Khalturina for assistance with RNAseq library preparation. This study was supported by funding provided by NSF EAPSI OISE-1311087, JSPS-SP-13027, PADI Foundation Grant 11199, Sigma Xi Grants-in-Aid of Research and Integrative Biology Department at Oregon State University to SAK. Corals were sampled under the Okinawa Prefecture permit No. 25-67.

## Data Accessibility

Raw RNAseq reads are available on the NCBI Sequence Read Archive https://www.ncbi.nlm.nih.gov/sra/PRJNA777284. Code to reproduce the analyses is available at: https://github.com/skitchen19/coral_larval_heatStress_colonization_expression.

## Author Contributions

SAK and VMW designed the experiment. SAK performed the experiment with field and laboratory support provided by SH, NS, and CS. SAK performed the data analysis and bioinformatics. DJ contributed statistical analysis on the survival data. SAK, NS, CS and VMW provided funding for the project. SAK, VMW, and CS wrote the manuscript and all authors contributed edits.

