## Supplemental Figures for "Coral larvae suppress the heat stress response during the onset of symbiosis thereby decreasing their odds of survival"

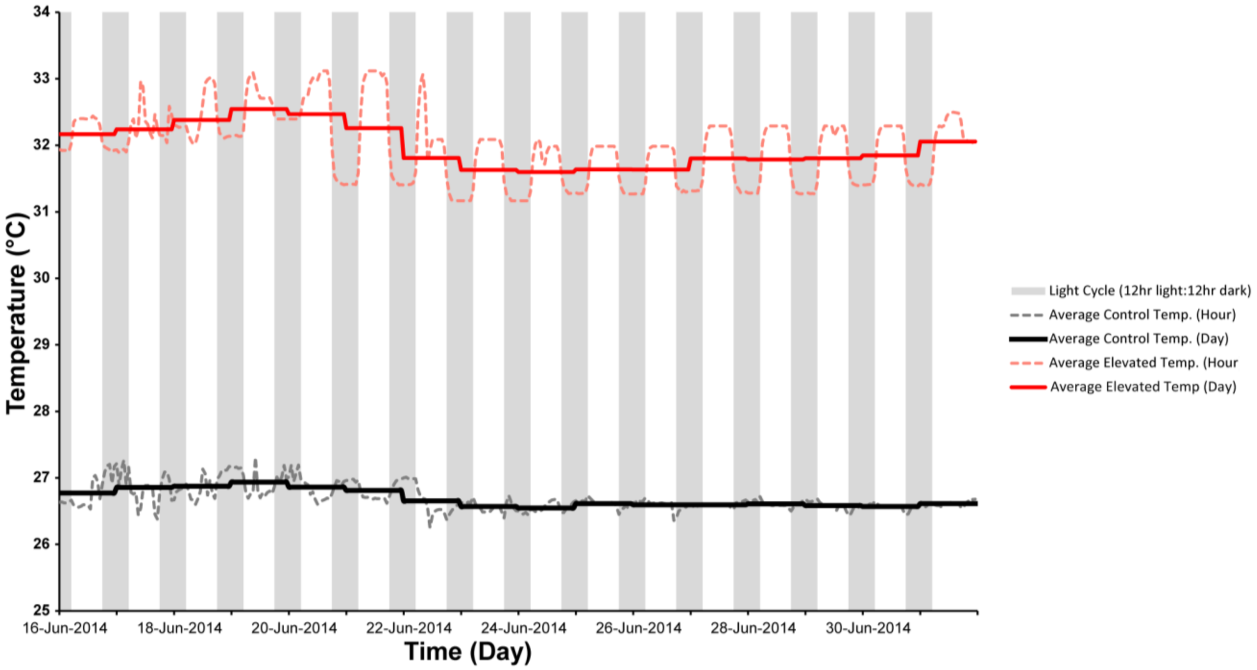


**Figure S1. Experimental temperatures over 14 days.** Temperature loggers submerged in water recorded temperature at 1 min intervals from each experimental chamber. Average temperatures by hour (dashed line) and by day (solid line) are displayed with the 12 h light: 12 h dark cycle indicated by shaded bars. Light cycle caused approximately 0.5 °C oscillation in temperature in the 32 °C treatment group, but not with control temperature. Control treatment = black; high temperature treatment = red.

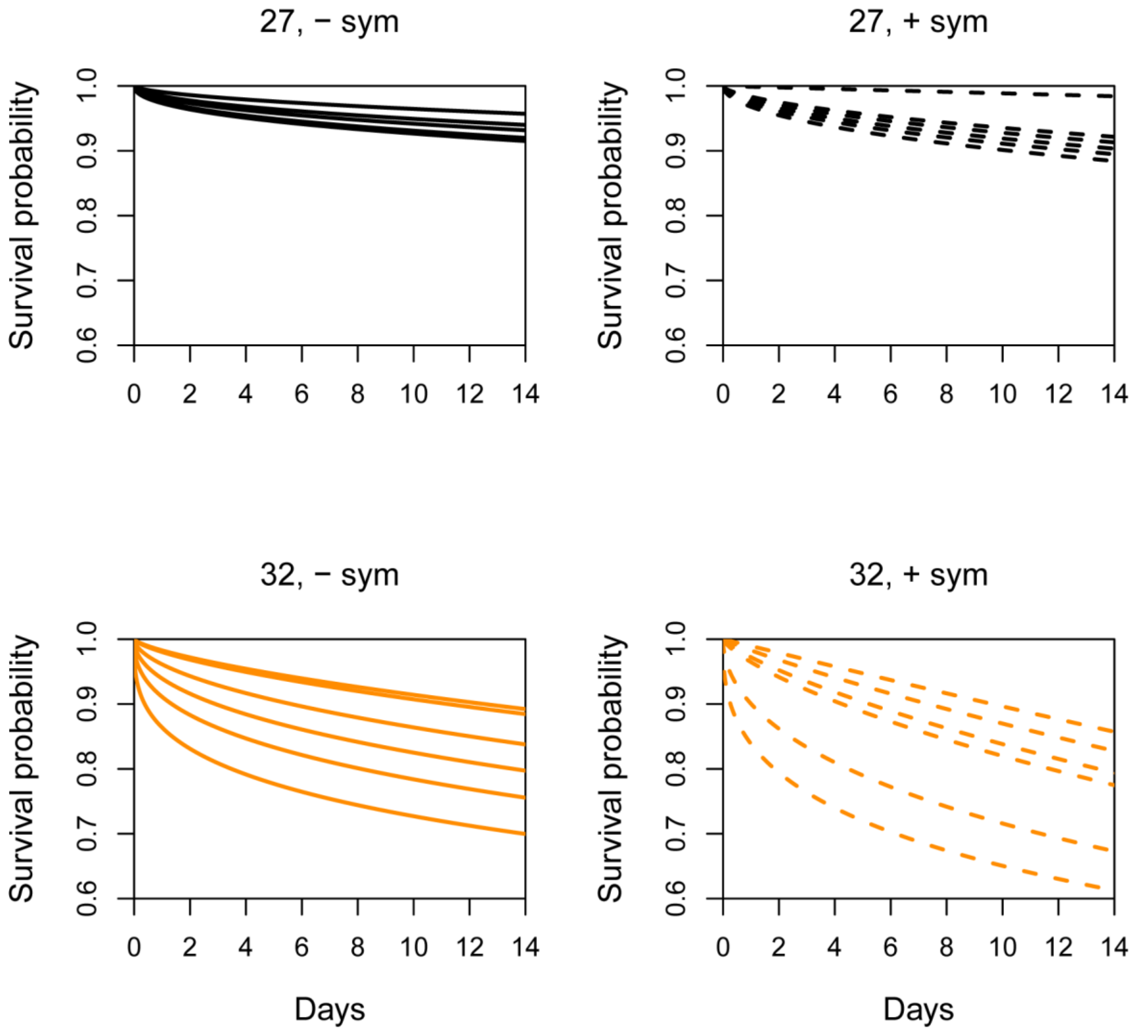


**Figure S2. Survival probability for each replicate tube in a given treatment.** Fitted survival probabilities under the Weibull proportional hazard model for each treatment group (*n*= 300 larvae), averaged across replicate tubes (*n*= 6 tubes per treatment group). The odds of survival decreased significantly with both temperature and symbiosis, but no interaction of these treatments was detected. Solid line = aposymbiotic (A), dashed line = symbiont colonized (S), black = 27 °C control temperature (C), orange = 32 °C high temperature (H).

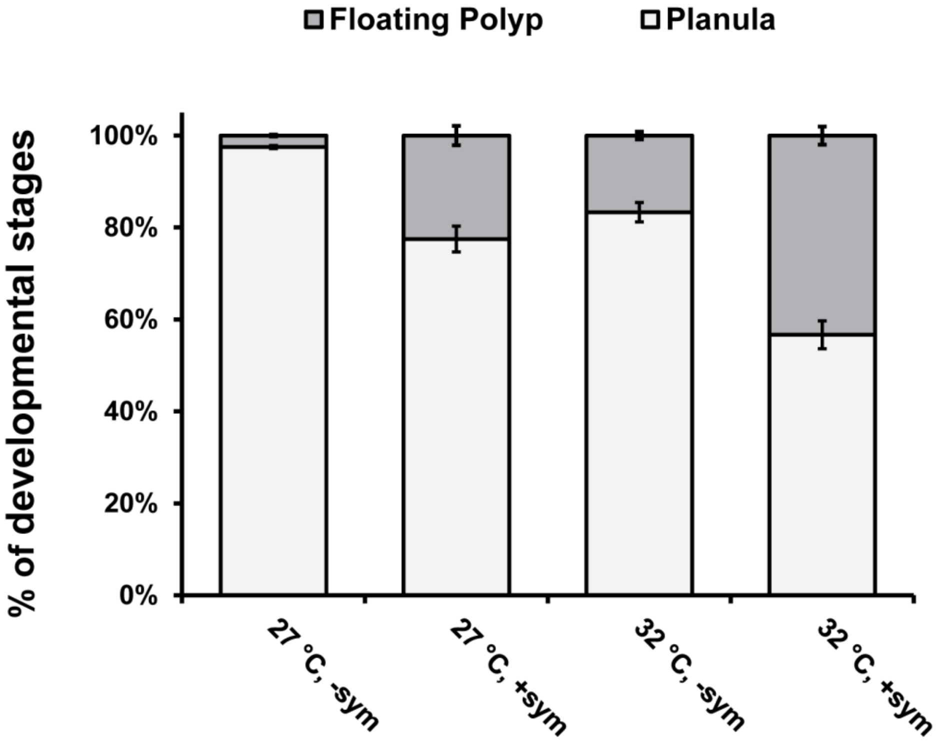


**Figure S3. Percentage of metamorphosed floating polyps at 3 dpi in samples used for RNAseq.** Twenty individuals were randomly collected from each replicate tube (*n*= 6 per group) and categorized into floating polyps (dark gray) or planulae (light gray). The odds of pre-mature development with temperature or colonization were tested using a logistic regression. The bars represent average for each developmental stage with standard error bars.

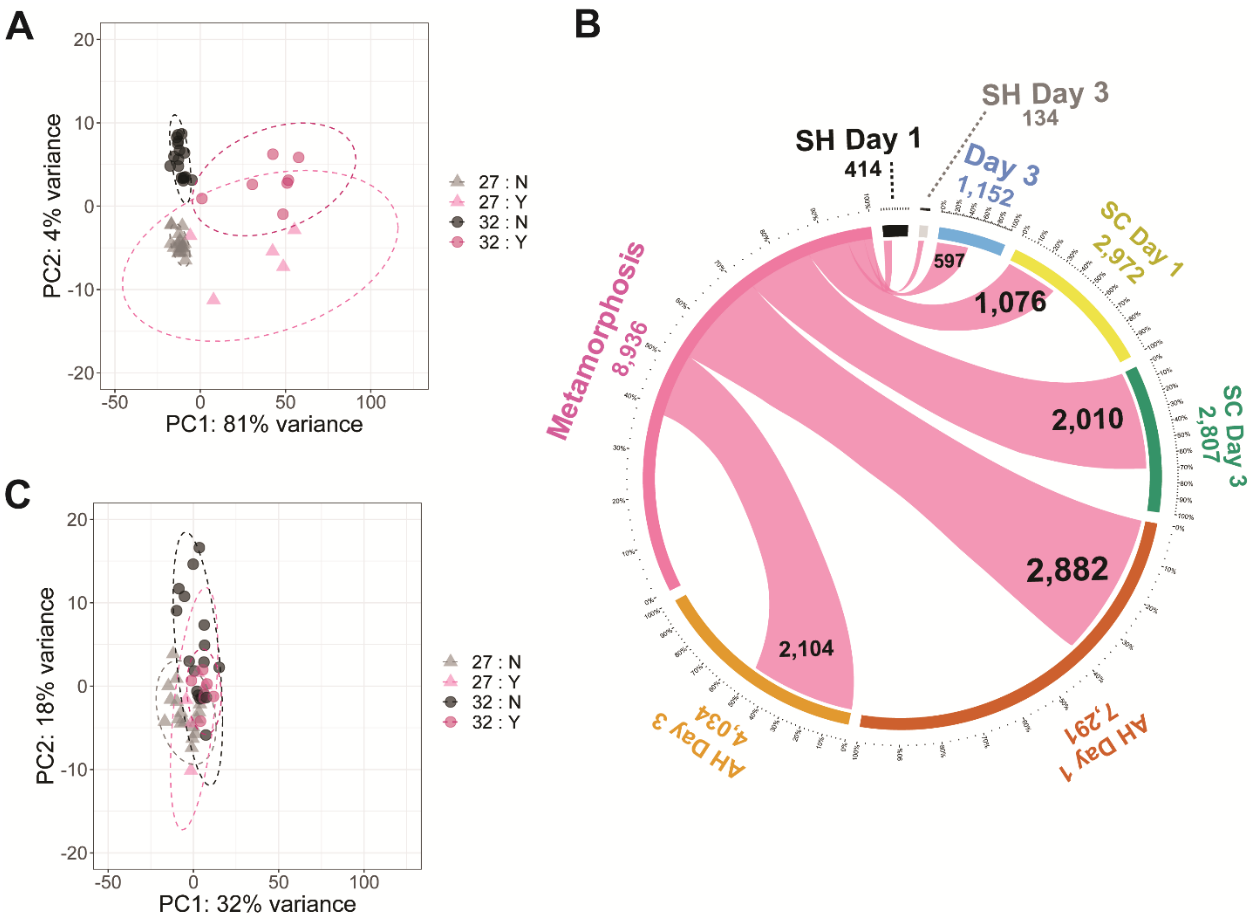


**Figure S4. Identification and removal of metamorphosis differentially expressed genes.** (A) Principle component analysis of top 500 genes with highest variance of rlog transformed counts for each *A. digitifera* larval sample. Larval samples that did not undergo metamorphosis (N) formed tight clusters with their respective temperature treatments (gray triangle = 27 °C, black circle = 32 °C). The samples that had some level of metamorphosis (Y) ranging from 5 to 85% did not fall within the non-metamorphosed groups, but were also separated by temperature (light pink triangle = 27 °C, dark pink triangle = 32 °C). (B) 8,936 DEGs identified with metamorphosis were shared with the respective treatments. The colored segments outside the circle are the total number of DEGs for each category. The inner pink segments show the number of those DEGs that are shared with metamorphosis. (C) Principle component analysis of rlog transformed counts after the removal of the metamorphosis DEGs.

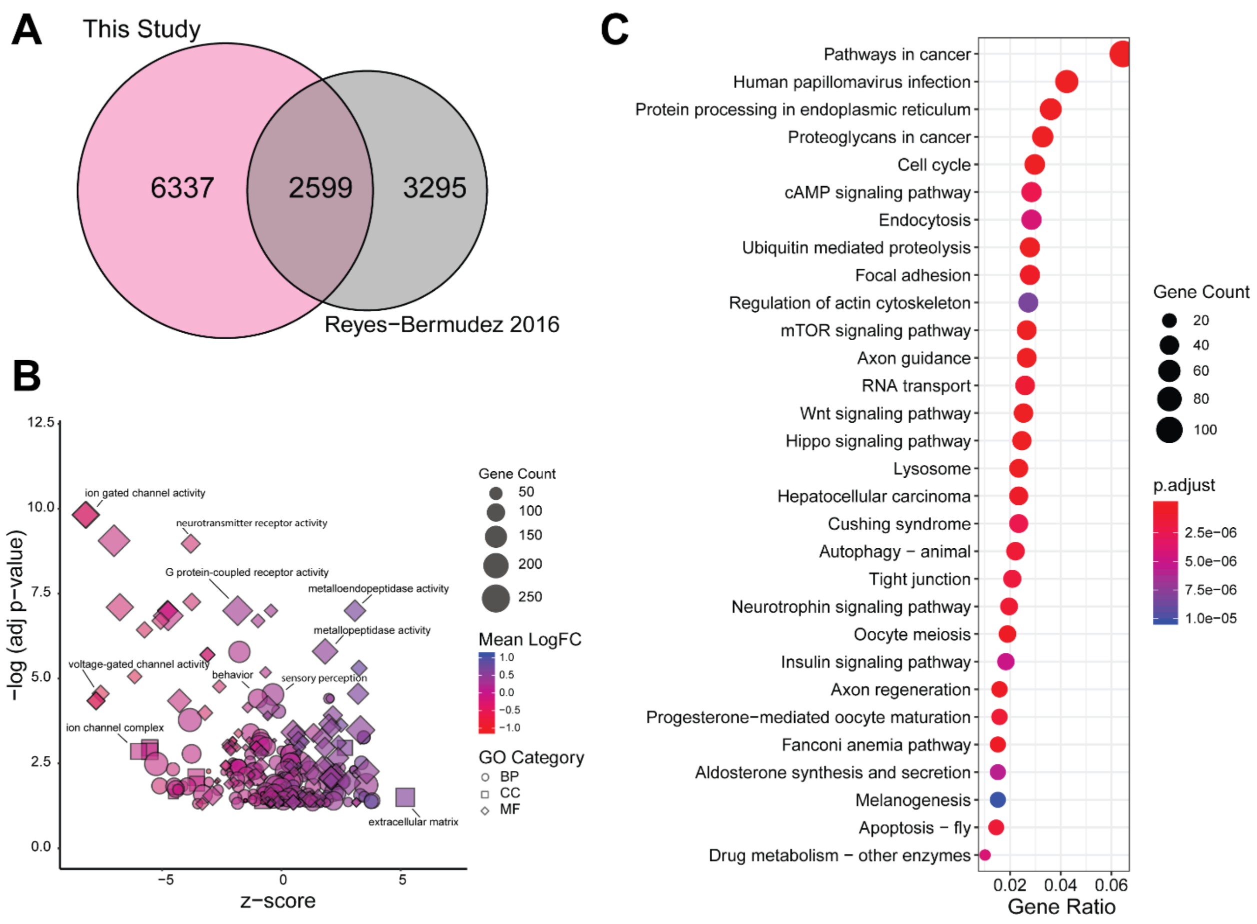


**Figure S5. Enrichment of GO terms and KEGG pathways in differentially expressed genes with metamorphosis.** (A) A Venn diagram shows overlap of 2,599 metamorphosis DEGs identified in this study with a previous study by Reyes-Bermudez et al. (2016) on the differences between planula larvae and adult *A. digitifera*. (B) Over-representation analysis identified 110, 159 and 16 GO terms from molecular function, biological process and cellular component categories, respectively. The adjusted p-values were $\log_{10}$ transformed for each term and a z-score was calculated based on the number of genes upregulated vs. downregulated out of all genes assigned to a given GO term. The shape is based on the GO category (shapes: MF= diamond, BP= circle, CC=square), the size is based on the number of genes, and the color is based on the mean $\log_2$ fold-change calculated for all genes for a given GO term. (C) KEGG pathway enrichment analysis identified 187 pathways, of which the top 30 are shown. The size of the circles is based on gene counts and the color is based on the adjusted p-value.

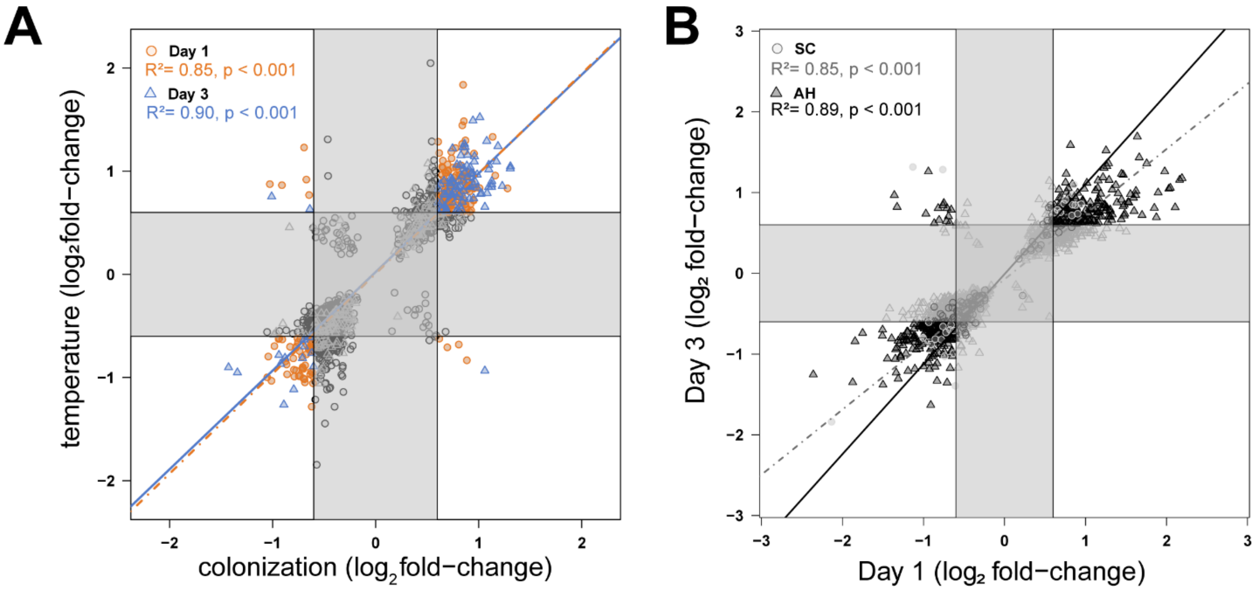


**Figure S6. Correlation of gene expression from SC- and AH-treated larvae.** (A) A positive correlation of DEGs shared between SC and AH treatment groups (n= 942 DEGs) on days 1 (orange circles) and 3 (blue triangles) (day 1 Pearson's correlation= 0.85, day 3 Pearson's correlation= 0.90). (B) Genes shared between days 1 and 3 by the SC (grey circles) or AH (black triangles) treatment groups were positively correlated (SC 1 and 3 dpi Pearson's correlation= 0.85, $p < 0.001$; AH 1 and 3 dpi Pearson's correlation = 0.890, $p < 0.001$). Genes within the gray boxes were differentially expressed but below the threshold of fold-change in this study, between 0.6 to -0.6.

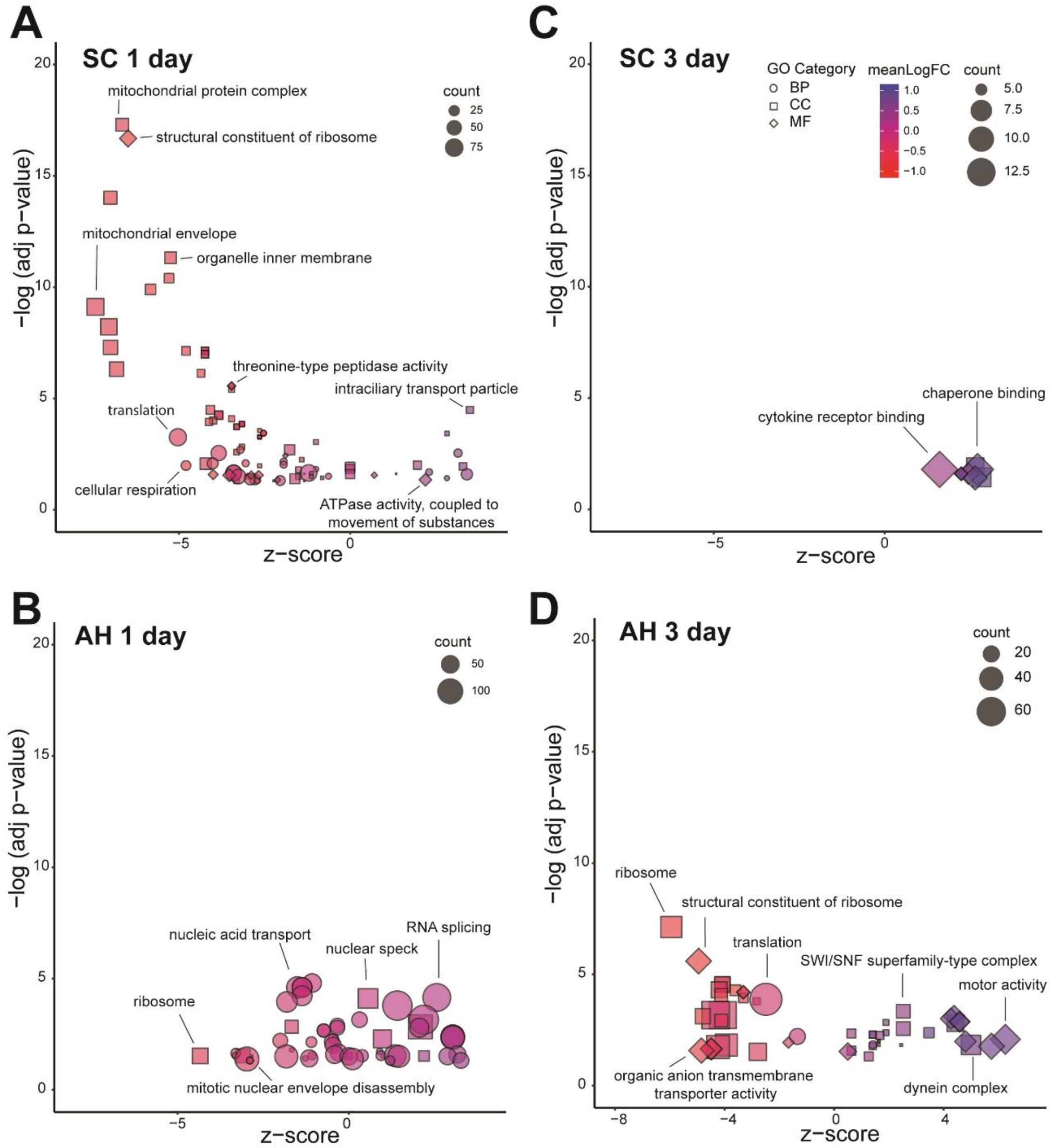


**Figure S7. Enriched GO terms for each treatment by day in *A. digitifera* larval samples.** GO term enrichment was performed on DEGs from day 1 SC (A) and AH (B) and day 3 SC (C) and AH (D) treatment groups with a hypergeometric model in clusterProfiler (Yu et al., 2012). The adjusted p-values were $\log_{10}$ transformed for each term and a z-score was calculated based on the number of genes upregulated vs. downregulated out of all genes assigned to a given GO term. The shape is based on the GO category (shapes: MF= diamond, BP= circle, CC=square), the size is based on the number of genes, and the color is based on the mean $\log_2$ fold-change calculated for all genes for a given GO term.

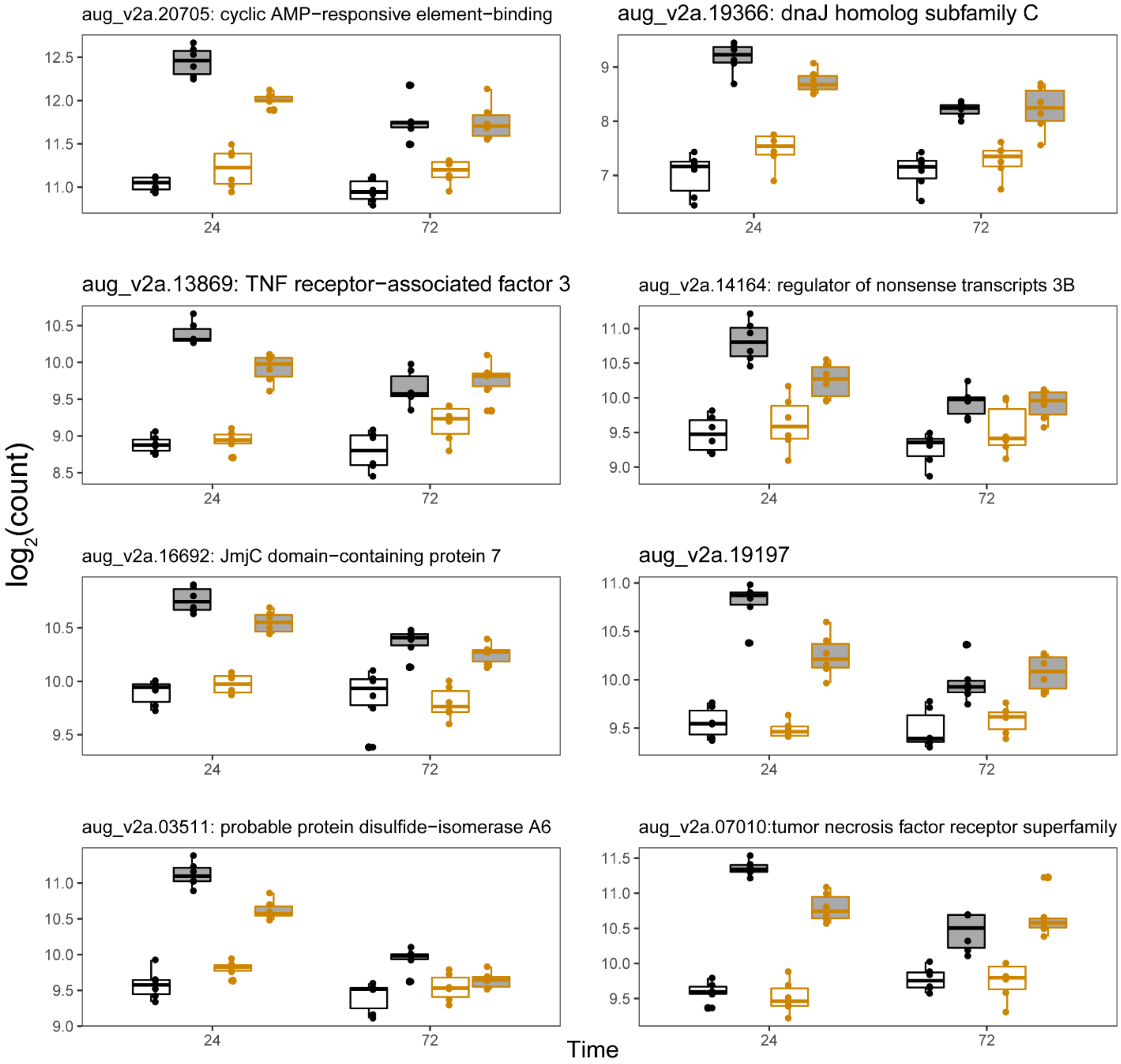


**Figure S8. Normalized gene expression of the top eight hub genes in module M21.** Black outline and white fill = AC, black outline and color file = AH, orange outline and white fill = SC, orange outline and color fill = SH.

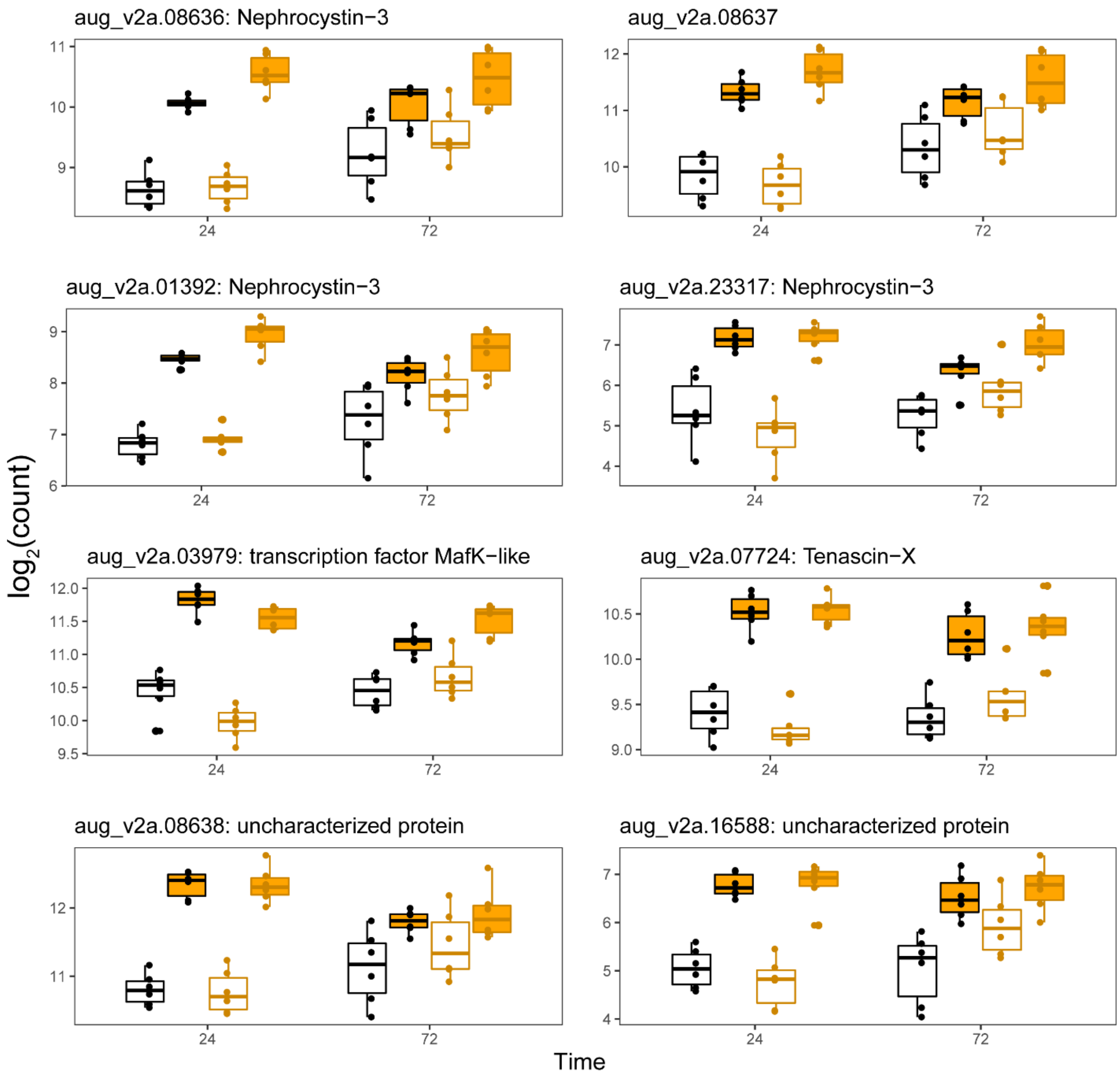


**Figure S9. Normalized gene expression of the top eight hub genes in module M12.** Black outline and white fill = AC, black outline and color file = AH, orange outline and white fill = SC, orange outline and color fill = SH.

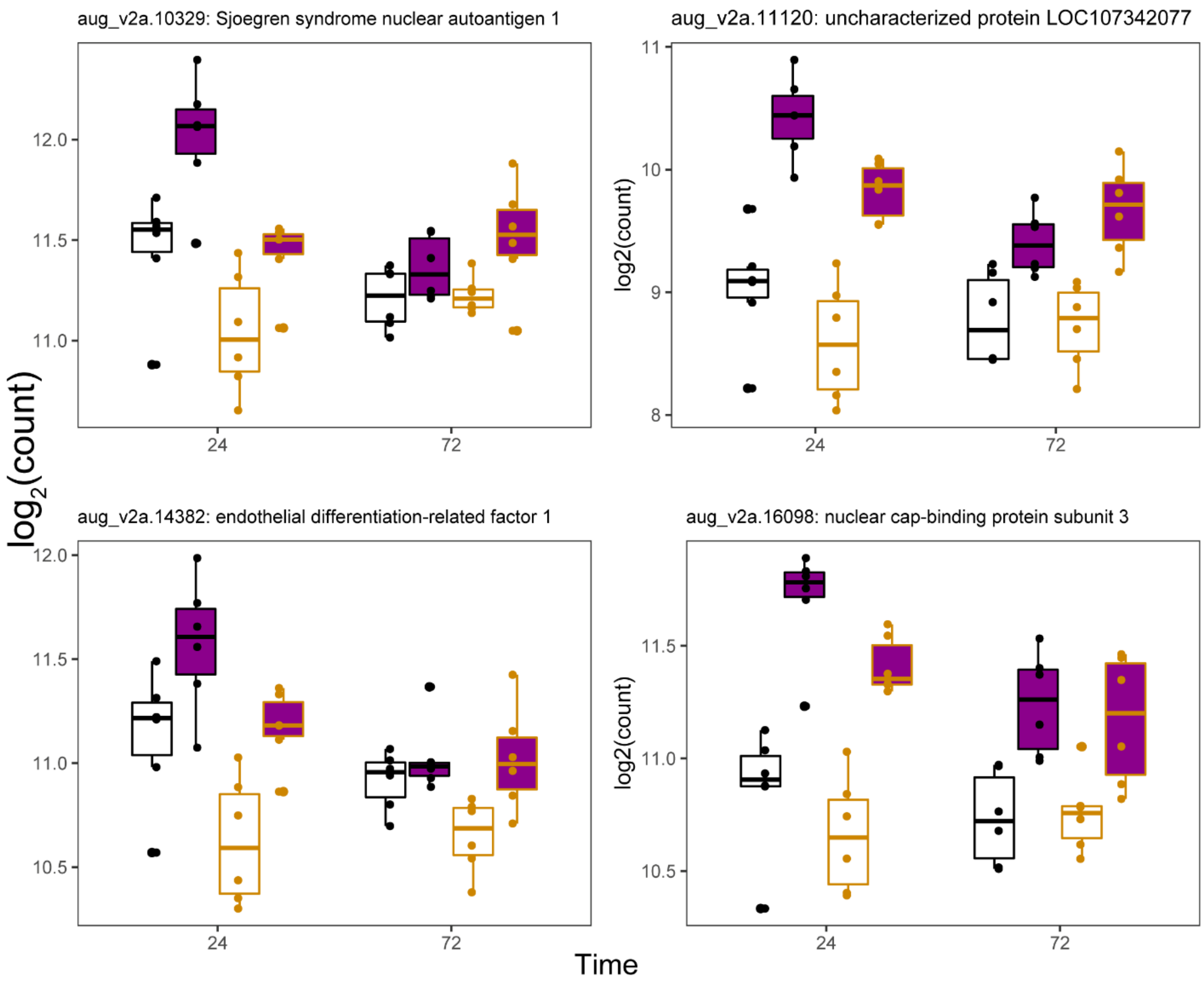


**Figure S10. Normalized gene expression of the top four hub genes in module M24.** Black outline and white fill = AC, black outline and color file = AH, orange outline and white fill = SC, orange outline and color fill = SH.

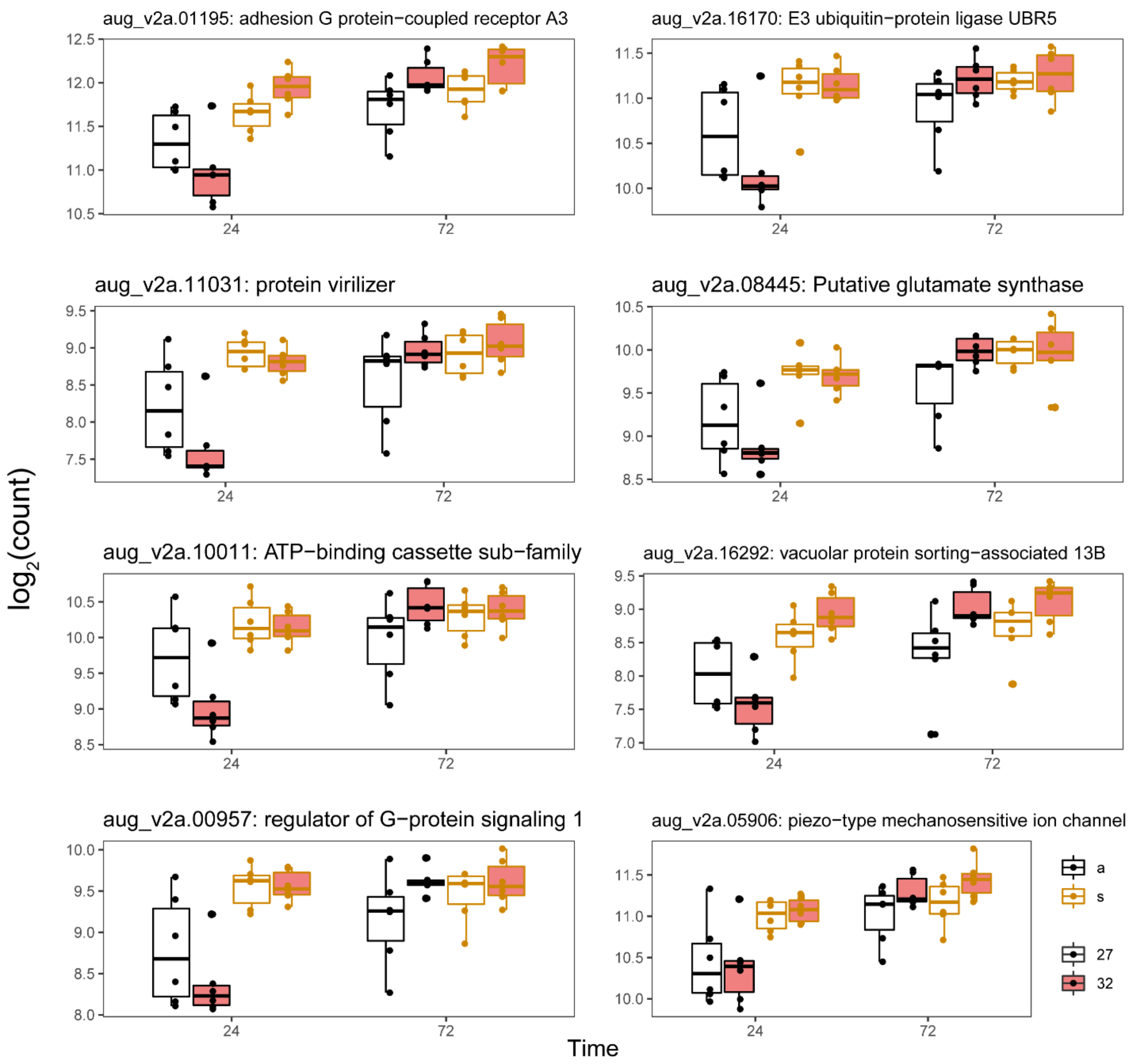


**Figure S11. Normalized gene expression of the top eight hub genes in module M13.** Black outline and white fill = AC, black outline and color file = AH, orange outline and white fill = SC, orange outline and color fill = SH.

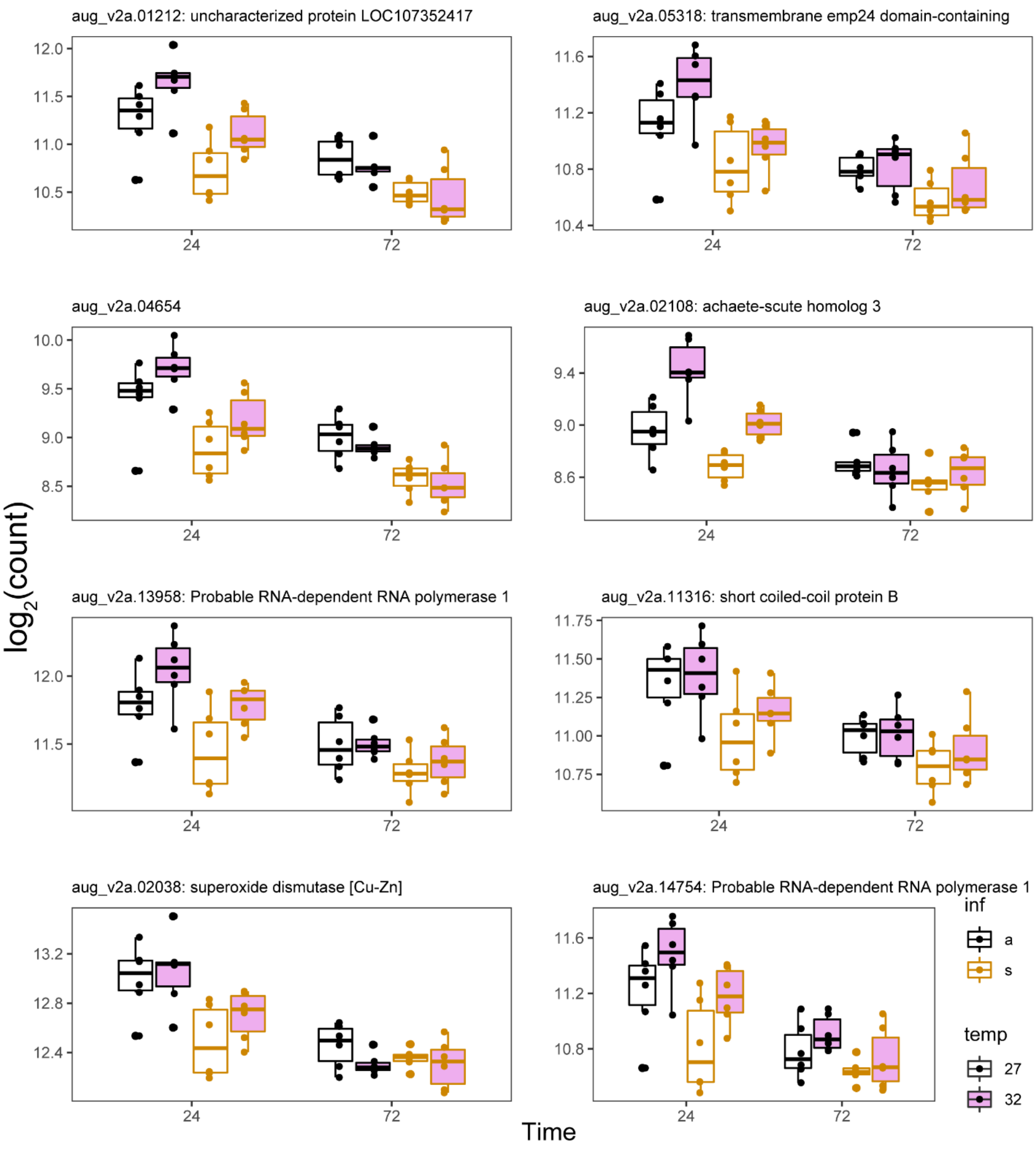


**Figure S12. Normalized gene expression of the top eight hub genes in module M25.** Black outline and white fill = AC, black outline and color file = AH, orange outline and white fill = SC, orange outline and color fill = SH.

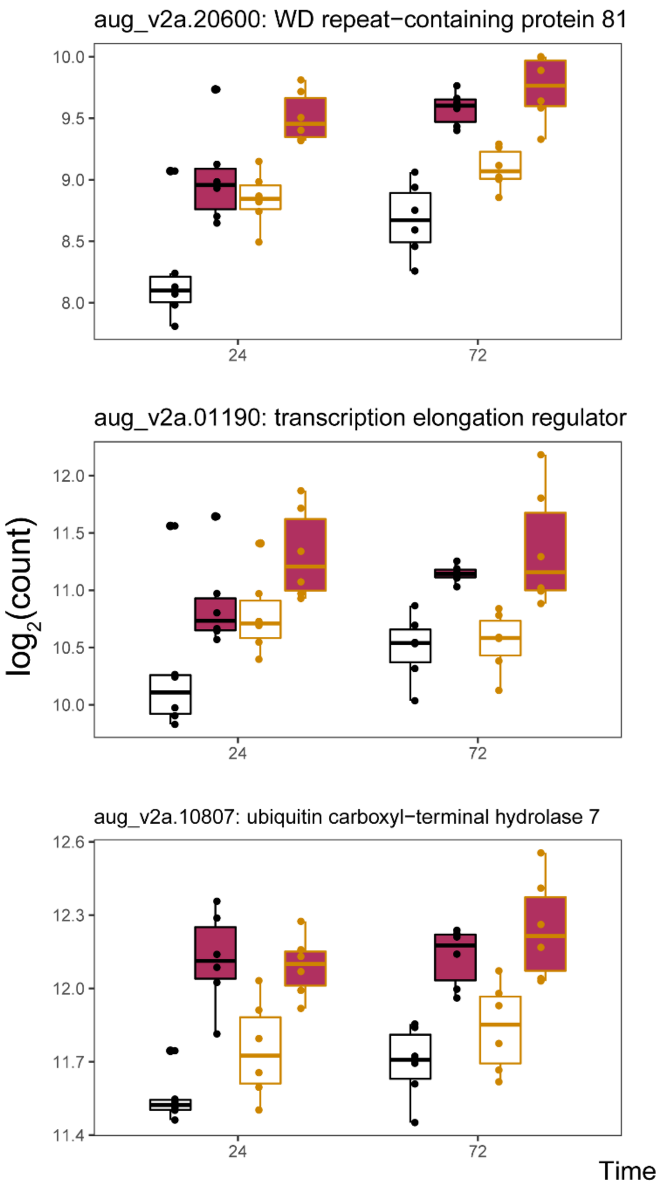


**Figure S13. Normalized gene expression of the five hub genes in module M14.** Black outline and white fill = AC, black outline and color file = AH, orange outline and white fill = SC, orange outline and color fill = SH.

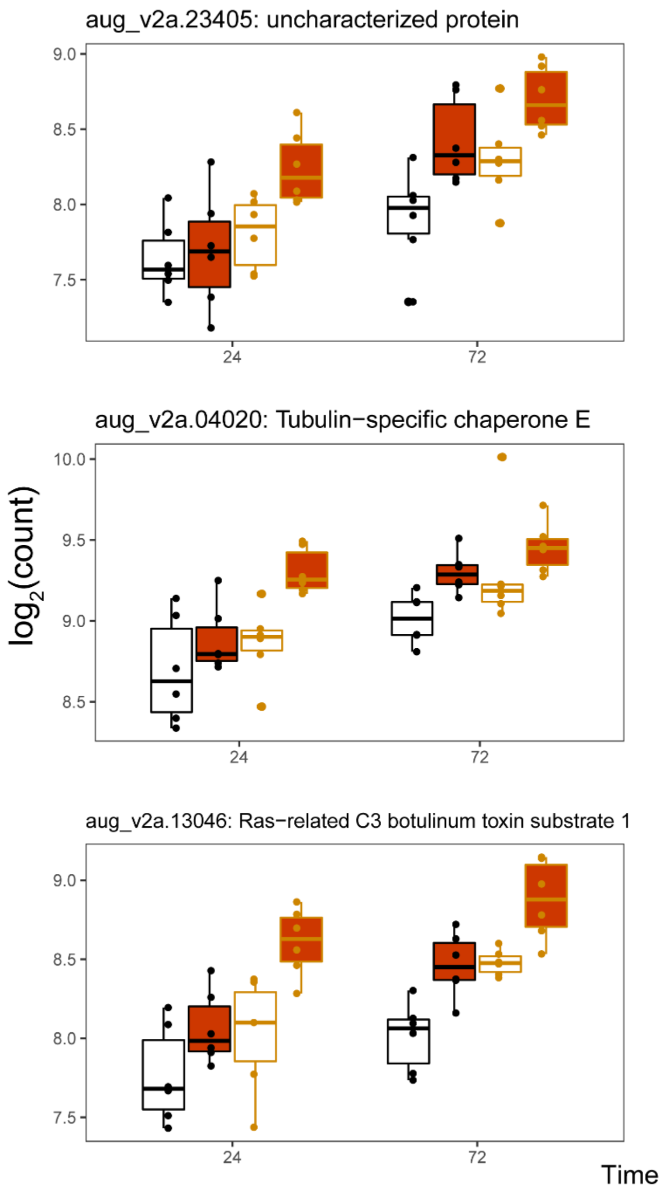


**Figure S14. Normalized gene expression of the five hub genes in module M15.** Black outline and white fill = AC, black outline and color file = AH, orange outline and white fill = SC, orange outline and color fill = SH.

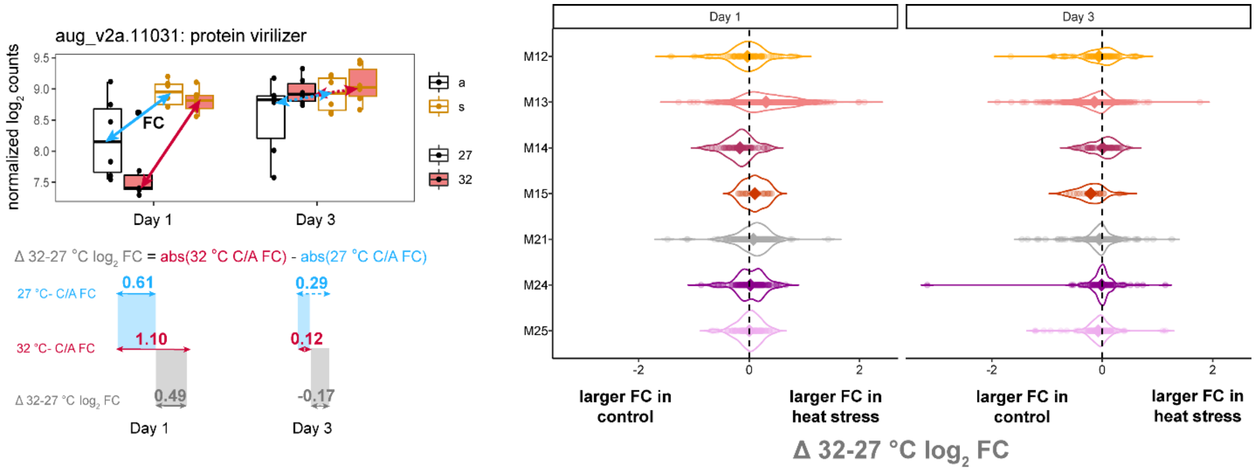


**Figure S15. Comparison of colonization fold-change in the elevated and control temperature treatments.** The difference in the absolute value of high temperature colonized larvae to not colonized larvae fold-change from the absolute value of the low temperature colonized larvae to not colonized larvae fold-change (Δ 32-27 °C $\log_2$ expression) was calculated as exemplified for one of the hub genes in module M13 that was highly correlated with colonization. Distribution of Δ 32-27 °C $\log_2$ expression for each significant gene (individual points) within each module is displayed by the violin plots with the mean difference represented by the diamond. Negative values indicate larger fold-changes in the control temperature treatment while positive values indicate larger fold-changes in the high temperature treatment.

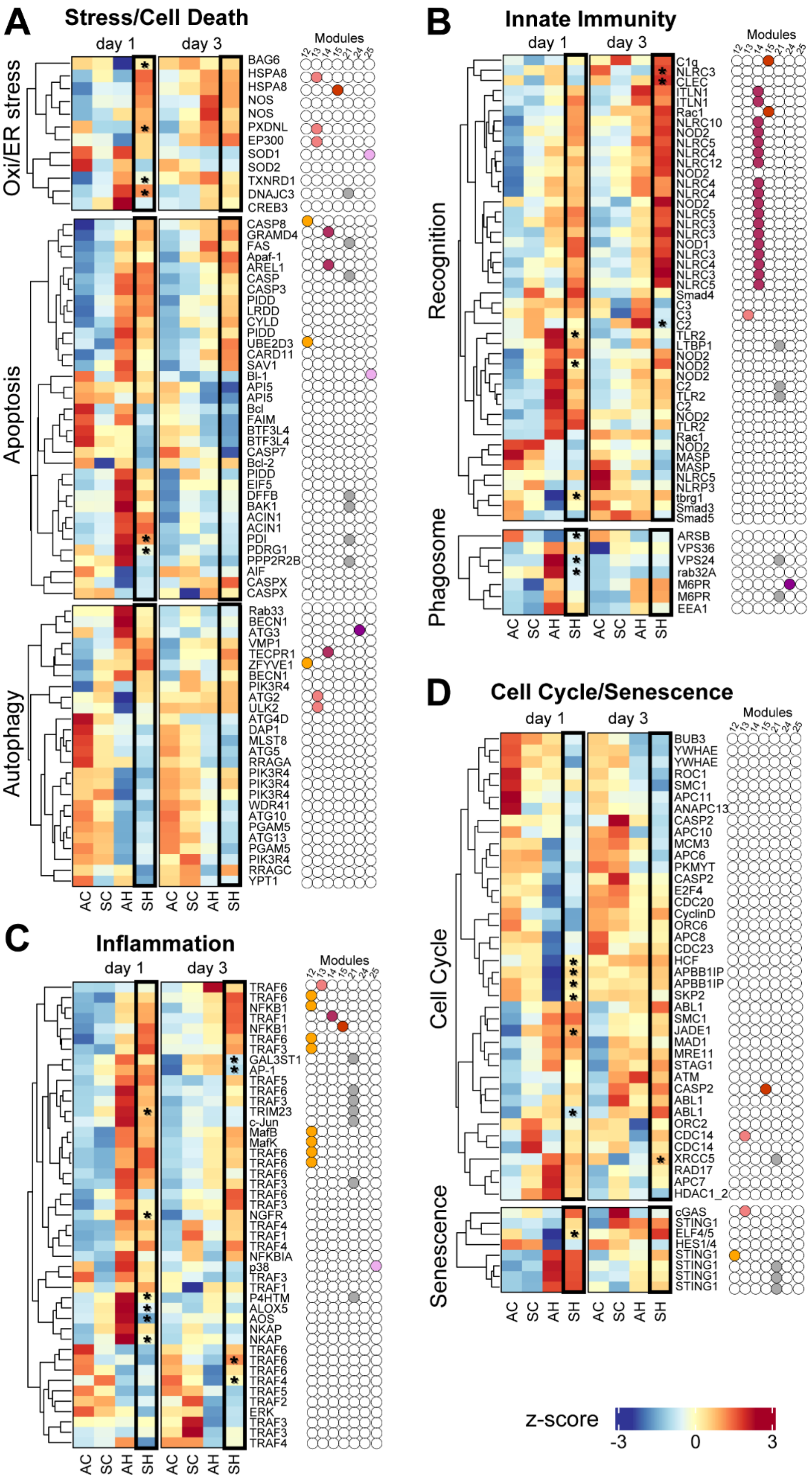
A
Stress/Cell Death
day 1
day 3
Modules
12 13 14 15 21 24 25
Oxi/ER stress
BAG6
HSPA8
HSPA8
NOS
NOS
PXDNL
EP300
SOD1
SOD2
TXNRD1
DNAJC3
CREB3
Apoptosis
CASP8
GRAMD4
FAS
Apaf-1
AREL1
CASP
CASP3
PIDD
LRDD
CYLD
PIDD
UBE2D3
CARD11
SAV1
BI-1
API5
API5
Bcl
FAIM
BTF3L4
BTF3L4
CASP7
Bcl-2
PIDD
EIF5
DFFB
BAK1
ACIN1
ACIN1
PDI
PDRG1
PPP2R2B
AIF
CASPX
CASPX
Autophagy
Rab33
BECN1
ATG3
VMP1
TECPR1
ZFYVE1
BECN1
PIK3R4
ATG2
ULK2
ATG4D
DAP1
MLST8
ATG5
RRAGA
PIK3R4
PIK3R4
PIK3R4
WDR41
ATG10
PGAM5
ATG13
PGAM5
PIK3R4
RRAGC
YPT1
AC SC AH SH AC SC AH SH
B
Innate Immunity
day 1
day 3
Modules
12 13 14 15 21 24 25
Recognition
C1q
NLRC3
CLEC
ITLN1
ITLN1
Rac1
NLRC10
NOD2
NLRC5
NLRC4
NLRC12
NOD2
NLRC4
NLRC4
NOD2
NLRC5
NLRC3
NLRC3
NOD1
NLRC3
NLRC4
NLRC3
NLRC5
Smad4
C3
C3
C2
TLR2
LTBP1
NOD2
NOD2
NOD2
C2
TLR2
C2
NOD2
TLR2
Rac1
NOD2
MASP
MASP
NLRC5
NLRP3
tbrg1
Smad3
Smad5
Phagosome
ARSB
VPS36
VPS24
rab32A
M6PR
M6PR
EEA1
AC SC AH SH AC SC AH SH
C
Inflammation
day 1
day 3
Modules
12 13 14 15 21 24 25
TRAF6
TRAF6
NFKB1
TRAF1
NFKB1
TRAF6
TRAF3
GAL3ST1
AP-1
TRAF5
TRAF6
TRAF3
TRIM23
c-Jun
MafB
MafK
TRAF6
TRAF6
TRAF6
TRAF3
TRAF6
TRAF3
NGFR
TRAF4
TRAF1
TRAF4
NFKBIA
p38
TRAF3
TRAF1
P4HTM
ALOX5
AOS
NKAP
NKAP
TRAF6
TRAF6
TRAF6
TRAF4
TRAF5
TRAF2
ERK
TRAF3
TRAF3
TRAF4
AC SC AH SH AC SC AH SH
D
Cell Cycle/Senescence
day 1
day 3
Modules
12 13 14 15 21 24 25
Cell Cycle
BUB3
YWHAE
YWHAE
ROC1
SMC1
APC11
ANAPC13
CASP2
APC10
MCM3
APC6
PKMYT
CASP2
E2F4
CDC20
CyclinD
ORC6
APC8
CDC23
HCF
APBB1IP
APBB1IP
SKP2
ABL1
SMC1
JADE1
MAD1
MRE11
STAG1
ATM
CASP2
ABL1
ABL1
ORC2
CDC14
CDC14
XRCC5
RAD17
APC7
HDAC1_2
Senescence
cGAS
STING1
ELF4/5
HES1/4
STING1
STING1
STING1
STING1
AC SC AH SH AC SC AH SH
z-score
-3
0
3

**Figure S16. Heat maps of the specific pathways differentially expressed in colonization and temperature treatments.** The average normalized, rlog counts of the transcripts (n=6 samples) involved in stress and apoptosis (A), immunity (B), inflammation (C), and cell cycle and cellular senescence (D) are presented. Expression was scaled row wise using a z-score transformation, ranging from -3 (blue) to 3 (red). Circles next to each heat map indicate the co-expression module membership of select modules referred to in the main text. Asterisk denote significant colonization-by-temperature interaction transcripts.
